# A system for inducible mitochondria-specific protein degradation *in vivo*

**DOI:** 10.1101/2022.12.20.521030

**Authors:** Swastika Sanyal, Anna Kouznetsova, Camilla Björkegren

**Affiliations:** Karolinska Institutet, Department of Biosciences and Nutrition, Neo, Hälsovägen 7c, 141 83 Huddinge, Sweden; Karolinska Institutet, Department of Cell and Molecular Biology, Biomedicum, Tomtebodavägen 16, 171 77 Stockholm, Sweden

## Abstract

Targeted protein degradation systems developed for eukaryotes employ cytoplasmic machineries to perform proteolysis. This has prevented mitochondria-specific analysis of genome maintaining proteins that localize to both mitochondria and nucleus. Here, we present an inducible mitochondria-specific protein degradation system in *Saccharomyces cerevisiae* based on the *Mesoplasma florum* Lon (mf-Lon) protease and its corresponding ssrA tag (called PDT). We show that mitochondrially targeted mf-Lon protease efficiently and selectively degrades a PDT-tagged reporter protein localized to the mitochondrial matrix. The degradation can be induced by depleting adenine from the medium and tuned by altering the promoter strength of the *MF-LON* gene. Finally, we demonstrate that mf-Lon degrades endogenous, dually localized proteins inside mitochondria. In summary, our system is an efficient tool for analysis of intricate mitochondria-nuclear crosstalk essential for proper mitochondrial function.

**One-Sentence Summary:** Mitochondria-specific protein degradation of dually localized proteins

---

Originating from alpha-proteobacteria, mitochondria have their own genome (*1, 2*). Approximately 1% of mitochondrial proteome is expressed by this genome, while the rest is nuclear-encoded, translated by cytoplasmic ribosomes, and guided by mitochondrial targeting signals (MTS) to their final location in mitochondria (*3, 4*). However, at least a third of the mitoproteome is localized to additional organelles (*5*). These dually localized proteins, for example those functioning in both the nucleus and mitochondria (*6*–*9*), generally score poorly for the features that define conventional MTSs (*10*), which prevents analysis of their mitochondria-specific functions by MTS disruption. Therefore, the development of a method for mitochondria-specific protein degradation is crucial.

To achieve this, we sought to engineer a system to degrade proteins specifically inside the mitochondria of living yeast cells. Given the prokaryotic origins of mitochondria, we borrowed from the so-called ribosome rescue mechanism, ubiquitously present in bacteria but absent from mitochondria (*11*). In this process, a degradation-inducing signal (called ssrA tag) is added at the end of a translating mRNA lacking a STOP codon. This allows release of the aberrant mRNA and subsequent proteolysis of the partially translated protein by endogenous proteases (*12*). We used the ssrA-degradation system from a simple mollicute, *Mesoplasma florum* (mf), which employs its Lon protease as the sole ssrA-degrading protease (*13*). We combined the mf-Lon protease with a variant of the mf-ssrA epitope called protein degradation tag [PDT, see Supplemental Text, and (*14*)], which allows a more faithful recognition by the mf-Lon protease (*14*).

To test if mf-Lon can perform PDT-degradation inside yeast mitochondria we created strains co-expressing mitochondrially targeted mf-Lon (mito-mf-Lon) and green fluorescent protein (GFP), either independently (mito-GFP), or tagged with PDT (mito-GFP-PDT) **(Fig. 1A)**. This revealed a significant loss of the GFP signal in the mitochondria of cells which expressed both mitochondrially-targeted GFP-PDT and mf-Lon, but not in the absence of mf-Lon, or when GFP lacked PDT or the MTS **(Fig. 1B-E)**. Cytoplasmically localized GFP-PDT signal was instead significantly stronger than its untagged counterpart **(Fig. 1D and E)**. The loss of mito-GFP-PDT signal in mito-mf-Lon expressing cells was robust, and more than 80% of the cells displayed an acute depletion of the signal **(fig. S1A and B)**. Together, this shows that mf-Lon can detect and specifically degrade PDT-tagged GFP inside mitochondria.

**Fig 1.**
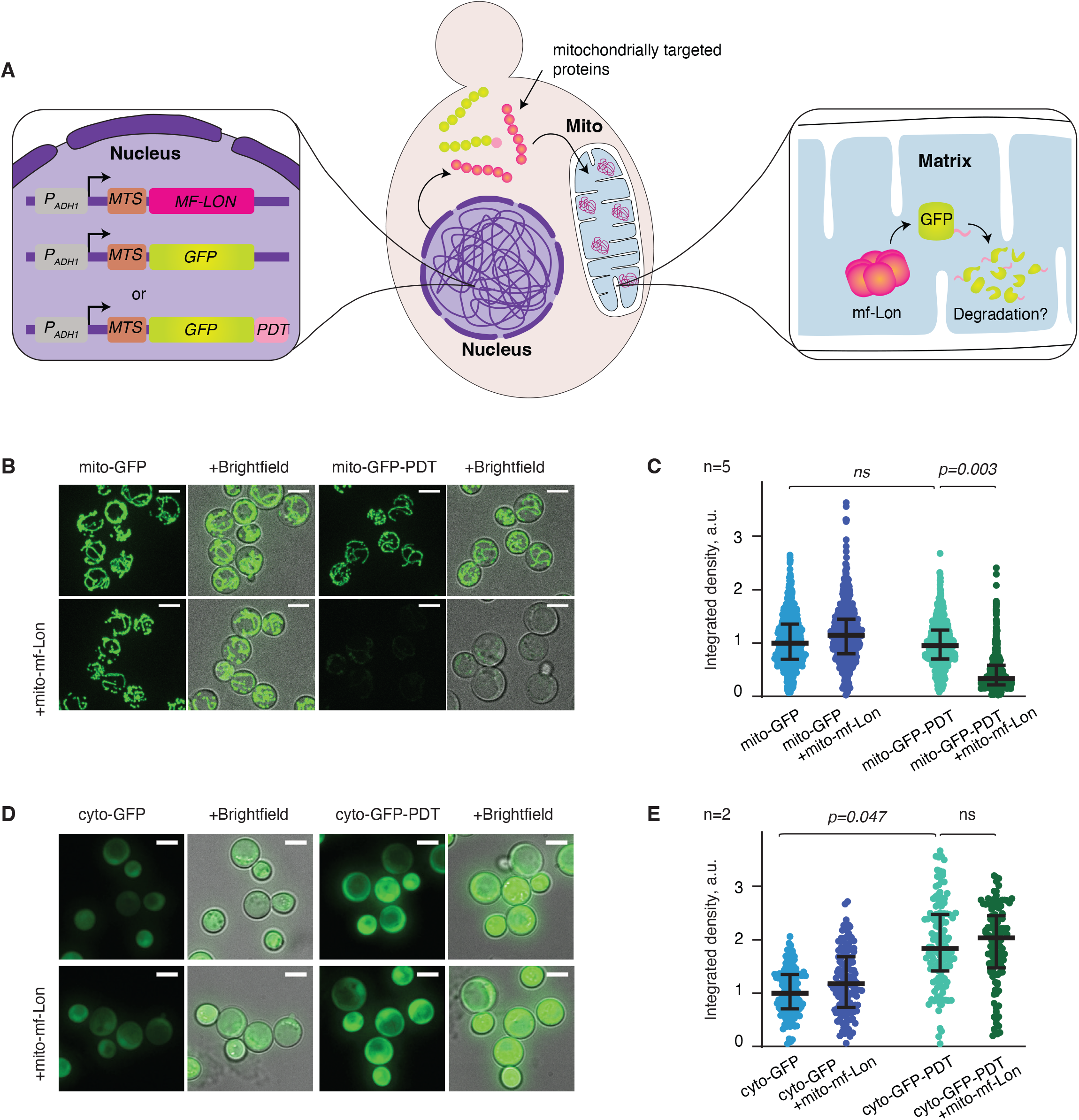
Mitochondria-specific degradation of GFP-PDT by *M. florum* Lon protease. **(A)** Schematics showing experimental set up. **(B)** Representative live cell images of yeast cells expressing mitochondrially-targeted GFP or GFP-PDT, either independently (upper panels) or together with mitochondrially-targeted *M*.*florum* Lon (lower panels). Cells were grown in standard minimal medium for ∼16-18 hours. Scale bar = 5 μm. **(C)** Dot plots showing quantification of GFP intensities as arbitrary units (a.u.) of cells indicated in (B). Each dot represents an individual cell. Data depict median value with interquartile range and were normalized to the mito-GFP strain. Pairwise comparisons were made by hierarchical resampling. P-value (*p*) more than 0.05 were considered not significant (*ns*). Five independent experiments were analyzed. **(D)** Similar to (B), but strains with GFP lacking the MTS are shown. **(E)** Similar to (C), except that dot plots show GFP intensities of cells (D), and data are normalized to the cyto-GFP strain.

Since respiration is dispensable in yeast, and the mitochondrial volume and metabolism changes during different growth phases and carbon sources (*15, 16*), we investigated if PDT degradation required specific growth conditions. We found that mf-Lon-induced mito-GFP-PDT degradation did not occur in undefined rich medium **(Fig. 2A)**. However, when a culture initiated in rich medium was transferred to synthetically defined minimal medium, the GFP-PDT signal was strongly reduced in mf-Lon expressing cells after overnight growth (16 h-20 h) **(Fig. 2B)**.

**Fig 2.**
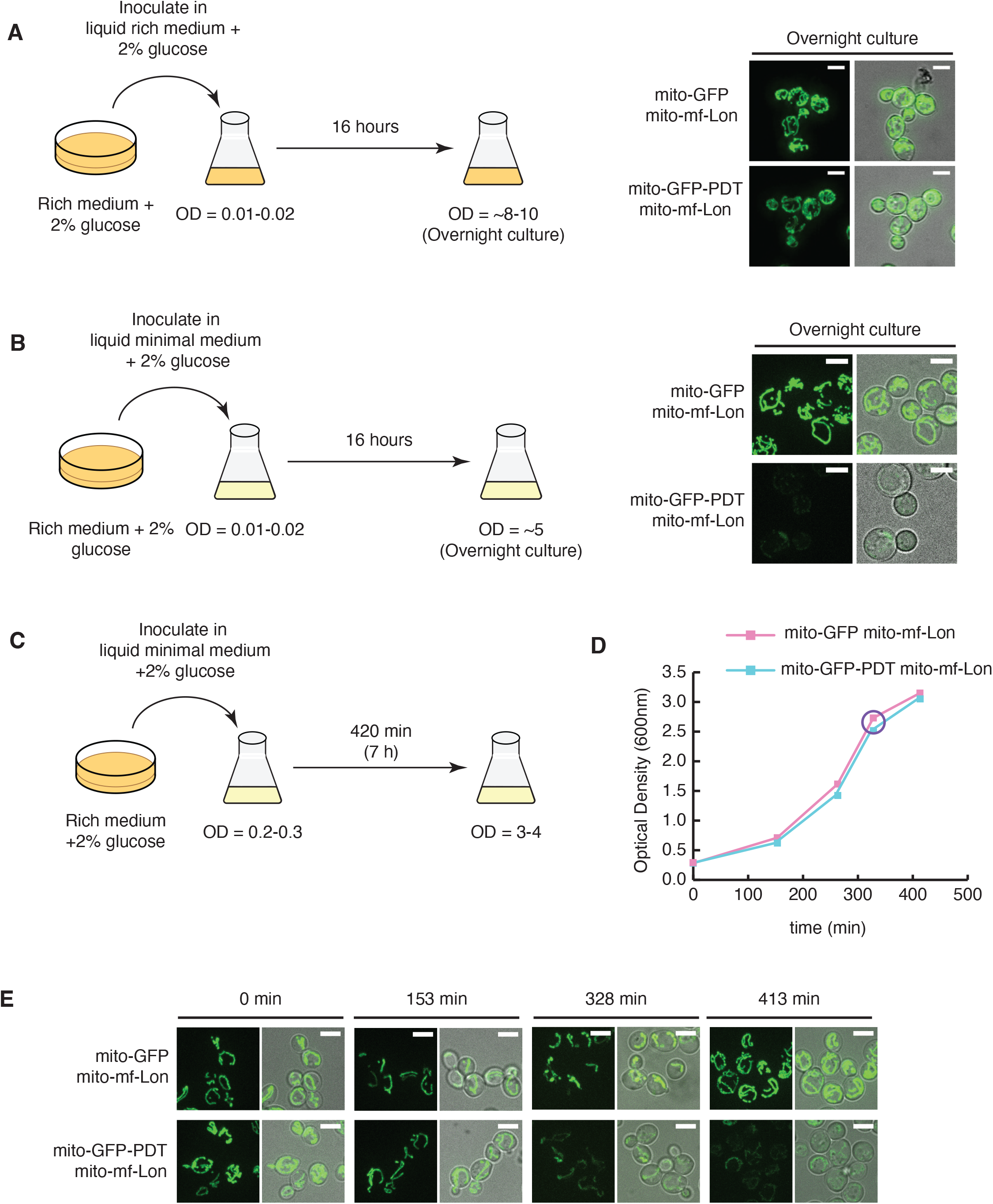
mf-Lon-induced PDT degradation is triggered at diauxie in minmal medium. **(A)** Cells were inoculated in rich dextrose medium and grown for 16 hours (schematic on left). Representative images of live cells (right). Scale bar = 5μm. **(B)** Cultures initiated in solid rich dextrose medium were grown in liquid minimal dextrose medium for 16 hours before being imaged (schematic on left). Representative images of live cells captured by confocal microscopy (right). Scale bar = 5μm. **(C-E)** Cells growing in solid rich medium were inoculated in minimal medium at indicated optical density (OD) **(C)** and analyzed for growth **(D**) and for GFP-PDT degradation **(E). (D)** Representative growth curve of cells grown in **(C)**. Purple circle indicates the time-point when GFP-PDT degradation was detected. **(E)** Live cell confocal images of samples taken at indicated time-points in **(D)**. Scale bar = 5μm.

To temporally determine when GFP-PDT was degraded during the overnight incubation period, we followed the GFP signal in mf-Lon cells co-expressing mito-GFP, or mito-GFP-PDT **(Fig. 2C)**. GFP-PDT degradation was initiated when cells entered deceleration growth phase, ∼5.5 hours after inoculation at an optical density (OD_600_) of 0.2, and continued to decrease during prolonged cell growth **(Fig. 2D and E)**. Contrarily, the signal of mitochondrial GFP lacking PDT increased during and after cellular growth slowed down **(Fig. 2E)**. These data demonstrate that the mitochondrial GFP-PDT degradation by mf-Lon occurs after cells have entered deceleration growth phase in synthetic minimal medium.

Yeast cell duplication decelerates when the glucose supporting rapid fermentative growth becomes limiting in the medium, causing cells to undergo a metabolic reprogramming that is needed for respiration of ethanol, which is the fermentation product of glucose (*15, 17*). This transition from fermentation to respiration is known as the diauxic shift and involves catabolite de-repression resulting from glucose exhaustion and is accompanied by an increase in mitochondrial DNA (mtDNA) (*18, 19*). Moreover, in yeast strains that are defective in adenine production, depletion of the base precedes glucose exhaustion, the diauxic shift is reached earlier, and growth retards even in the presence of sufficient extracellular glucose (*20*).

Taking advantage of the fact that the strain used for the investigation is an adenine auxotroph, we tested whether adenine can be used to control GFP-PDT degradation. Supplementing standard minimal medium with 50 mg/l adenine prevented GFP-PDT degradation **(fig. S2A)**, establishing that adenine depletion is central to the degradation in post-diauxic cells. To test this directly, we resuspended cells growing in an excess of adenine in medium containing 0, 2, or 20 mg/l adenine, and at a glucose concentration of 1%, which is similar to extracellular glucose concentration when cells reach diauxie (*15, 20*) **(fig. S2B)**. The mitochondrial GFP-PDT signal in mf-Lon expressing cells was reduced after 2-3 hours of adenine withdrawal (0 mg/l) or limitation (2 mg/l), and was further reduced during continued growth without adenine. Contrarily, the signal of GFP lacking PDT significantly increased in intensity under the same conditions, despite the presence of mf-Lon **(Fig. 3A-C)**. Likely, this increase reflects the reorganization and enlargement of the mitochondrial network and volume, a characteristic of post-diauxic cells (*15, 17*). Together, this establishes that mf-Lon-induced mito-GFP-PDT degradation can be triggered by adenine depletion, thereby providing an inducible system for mitochondria-specific protein degradation.

**Fig 3.**
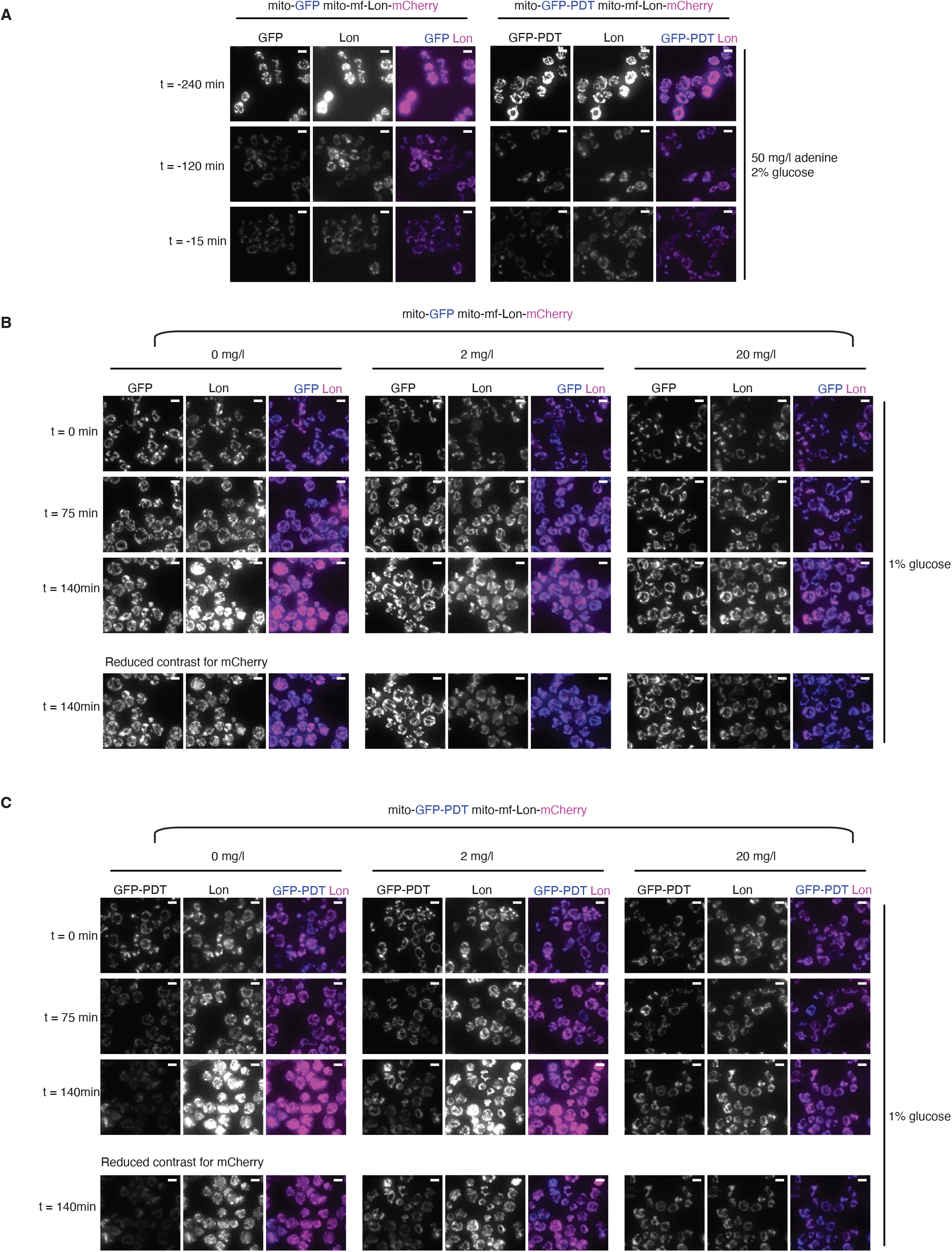
Adenine limitation induces PDT degradation by mf-Lon protease. **(A)** Live cell images of indicated cells growing in minimal media supplemented with 2% glucose and 50 mg/l adenine, before they were shifted to media containing 1% glucose and 0, 2, or 20 mg/L adenine. See fig. S2B for schematics. **(B)** Live cell images of mito-GFP mito-mf-Lon-mCherry cells growing in media containing 1% glucose and 0, 2, or 20 mg/l adenine at indicated time points. **(C)** Same as (B) but with mito-GFP-PDT mito-mf-Lon-mCherry cells. **(B and C, bottom panel)** Reduced contrast images of indicated time-point to diminish a background signal in the channel for mCherry recording, caused by a red pigment (oxidized ribosylaminoimidazole) accumulating in the vacuole upon adenine depletion. Scale bar = 5 µm.

High levels of Lon protease have been shown to downregulate mitochondrial DNA content (*21*). As we used the strong *ADH1* promoter (*P*_*ADH1*_) for the expression of our constructs **(Fig. 1A)**, we next performed Sybr Green I staining to analyse the number, morphology and overall distribution of mtDNA nucleoids **(fig. S3A)** (*22*). We found that mf-Lon expression from the *ADH1* promoter left total staining intensity per cell unchanged, implying that total mtDNA content was unaltered as compared to wild type **(fig. S3B)**. This was corroborated by mtDNA copy number measurement by quantitative PCR (qPCR) after induction of PDT degradation **(fig. S3G)**. However, the total number of detected mtDNA spots per cell, representing the number of nucleoids, was decreased **(fig. S3C)**. Moreover, total area occupied by mtDNA was decreased **(fig. S3D)**, and the intensity and area of each nucleoid increased, as compared to wild type cells **(fig. S3E and F)**. Together this shows that mf-Lon expression from the *ADH1* promoter does not affect the level of mtDNA content, but reduces nucleiod number and changes their morphology, possibly increasing their level of compaction.

To circumvent this effect, we sought to improve the system, aiming for a level of Lon protease closer to that of wild type. Given that the yeast endogenous Lon, Pim1, did not contribute to the GFP-PDT degradation, and mf-Lon expressed from *P*_*ADH1*_ could perform Pim1 respiratory function **(fig. S4A and C)** (*23*), we altered the levels of mf-Lon by replacing the *PIM1* ORF with the coding sequences for MTS-mf-Lon (*pim1Δ::MITO-MF-LON*). Similar to mf-Lon expressed from *ADH1* promoter, mf-Lon expressed from the *PIM1* locus rescued the respiratory defect of *pim1Δ* cells **(fig. S4A-B)**. Importantly, mitochondrial GFP-PDT was also degraded in *PIM1-*replaced*-*mf-Lon expressing cells, albeit at a somewhat lower efficiency, in line with reduced mf-Lon expression **(fig. S4D)**. Nucleoid staining also showed that even if total mtDNA content increased in the *PIM1-*replaced mf-Lon expressing cells **(fig. S3B and G)**, the number of nucleoids and total area occupied by mtDNA remained at wild type levels **(fig. S3C and D)**. Moreover, even though individual nucleoids stained more intensively in *PIM1-*replaced mf-Lon cells, they were significantly weaker than in cells that expressed mf-Lon from *P*_*ADH1*_ promoter **(fig. S3E and F)**. Together these data indicate that mf-Lon expression from the weaker *PIM1* promoter only has a mild effect on mitochondrial nucleoid number and morphology as compared to *P*_*ADH1*_-controlled expression.

Finally, in a proof-of-concept experiment, we tested whether mf-Lon degrades PDT-tagged endogenous mitochondrial proteins. We analyzed two proteins which localize both to the nucleus and mitochondria: the abasic endonuclease Apn1, and the helicase Pif1 (*24, 25*). We PDT-tagged Apn1 and Pif1 in cells expressing mf-Lon either from the *ADH1* promoter (*P*_*ADH1*_-mito-mf-Lon) or the *PIM1* locus (*pim1Δ::*mito*-*mf-Lon), and grew them under the conditions that induce mito-GFP-PDT degradation **(fig. S5A and B)**.

Western Blot analysis of whole cell extracts revealed that the levels of Apn1-PDT were similar in uninduced or induced *P*_*ADH1*_-mf-Lon expressing cells **(fig. S5C)**. In contrast, the Apn1-PDT levels were significantly reduced in crude preparations of mitochondria from the induced mf-Lon expressing cells **(fig. S5C and D, Fig. 4A)**. This degradation was even more evident after additional proteinase K treatment, which removes nuclear contaminations **(fig. S5C)**.

**Fig 4.**
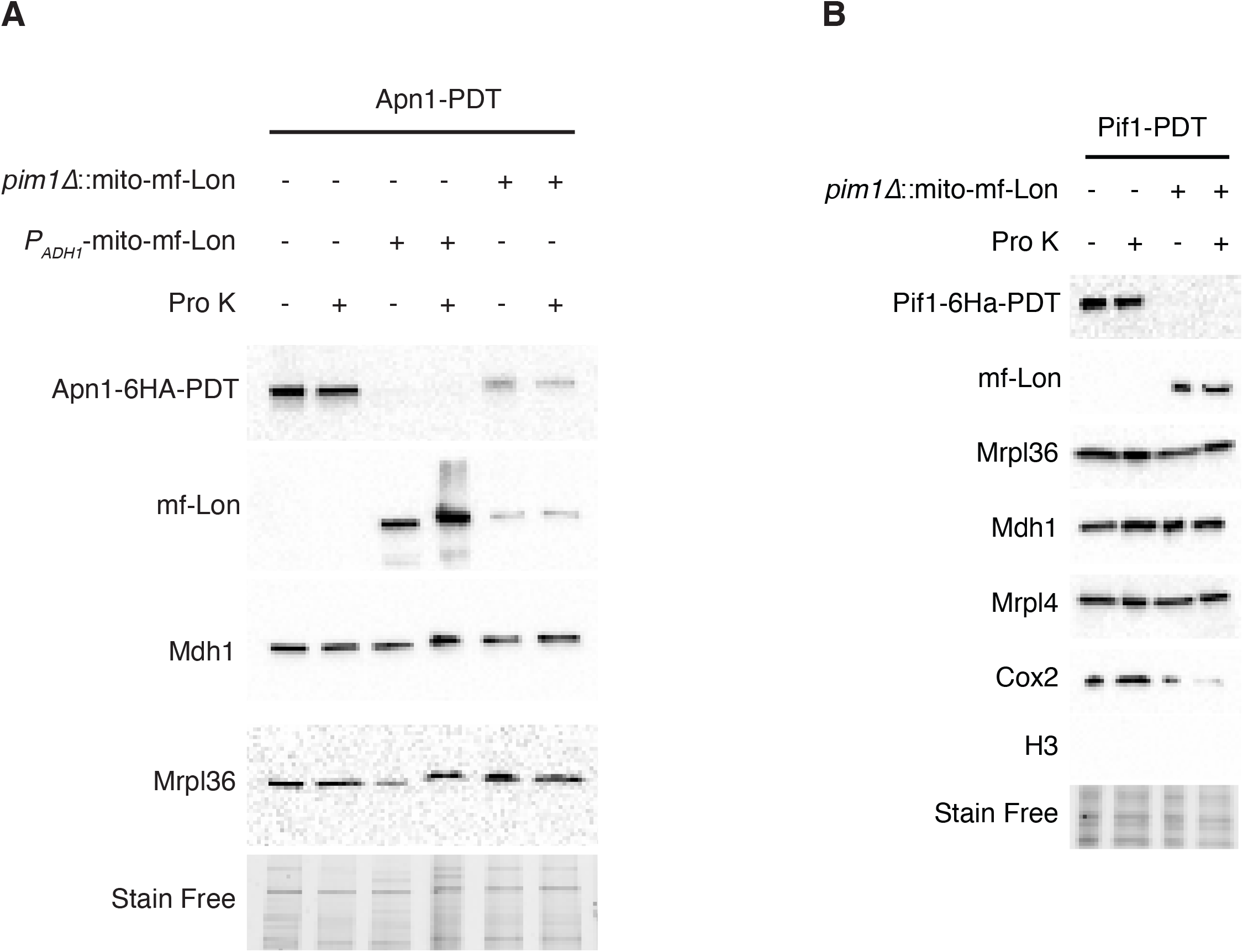
mf-Lon protease degrades yeast endogenous mitochondrial proteins tagged with PDT. **(A)** Western blot analysis of the mitochondria isolated from Apn1-PDT cells expressing mf-Lon from *ADH1* promoter (*P*_*ADH1*_-mito-mf-Lon) or the *PIM1* locus (*pim1Δ::*mito-mf-Lon) after induction of PDT degradation. Samples were treated with Proteinase K to remove extra-mitochondrial contamination. See supplementary figure 5D for additional markers. **(B)** Western blot analysis of mitochondria isolated from Pif1-PDT cells expressing *pim1Δ*::mf-Lon after induction of PDT degradation, and treated with Proteinase K. Stain free image serves as a loading control. The sizes of the proteins are: Apn1-PDT (anti-HA), ∼60 kDa; Pif1-PDT (anti-HA), ∼100 kDa; H3 (anti-H3), ∼15 kDa; Cox2 (anti-MTCO2), ∼35 kDa, Mrpl4 (anti-Mrpl4), ∼44 kDa; mf-Lon (anti-Flag), ∼100 kDa; Mdh1 (anti-Mdh1), 35.6 kDa; and Mrpl36 (anti-Mrpl36), ∼20 kDa.

Analysis of the *PIM1-*replaced mf-Lon strain also revealed mitochondria-specific degradation of Apn1-PDT, but at lower levels, in line with the reduction of promoter-strength **(Fig 4A, fig. S5D)**. Similarly, mf-Lon expression from *PIM1* locus was sufficient for complete degradation of the lesser expressed Pif1-PDT **(Fig. 4B)**. Importantly, except for a modest reduction of the respiratory Complex IV subunit, Cox2, a wide variety of mitochondrial matrix proteins, including mitochondrial ribosomes, were unaffected by mf-Lon expression **(Fig. 4B, fig. S5C and D)**. Overall, these data demonstrate that the mf-Lon can efficiently degrade yeast endogenous mitochondrial proteins in a promoter-strength dependent manner.

In conclusion, using the *Mesoplasma florum* Lon protease and its corresponding degradation tag, we show that mitochondria-specific protein degradation in living yeast cells is possible, can be induced through adenine depletion, and tuned through alterations in the expression levels of *MF-LON* and adenine concentration. The conditional degradation of protein of interest in mitochondria is especially powerful to investigate the role of proteins with essential functions. Furthermore, re-addition of adenine to cultures induced for PDT degradation of a protein of interest should restore its levels, thereby providing additional information on its molecular function. Together with recent advances on gene silencing in isolated mitochondria (*26*), our *in vivo* approach for targeted mitochondrial protein degradation significantly expands the molecular toolbox for analysis of mitochondrial functions of nuclear-encoded mitoproteins. This, in turn, will provide new insights on mitochondrial dysfunction which is fundamental to neurodegenerative disorders and ageing.

## Acknowledgments

We are thankful to Maria Falkenberg, Martin Ott and the members of CB lab for helpful discussions. We thank Martin Ott for antibodies for mitochondrial matrix markers. This study was funded by a post-doctoral grant from Wenner-Gren Foundations to SS, a Swedish Research Council grant (DNR 2021-01118) to AK, and a Knut and Alice Wallenberg foundation project grant to CB. Microscopy was performed at the Live Cell Imaging Core facility/Nikon Center of Excellence, at Karolinska Institutet, supported by the KI infrastructure council.

## Author contributions

Conceptualization: SS and CB

Methodology: SS and CB

Investigation: SS

Visualization: SS and AK

Funding acquisition: CB

Project administration: CB

Supervision: CB

Writing – original draft: SS

Writing – review & editing: SS, AK and CB

## Data and materials availability

All materials used in the analysis are available upon request to corresponding authors. All data are available in the main text or the supplementary materials.

## Supplementary Materials

Materials and Methods

Supplementary Text

Figs. S1 to S6

Tables S1 to S2

References (*27*–*41*)

## Materials and Methods

### Construction of plasmids

For mitochondrially targeted GFP, we used the pVT100U-mtGFP (Addgene) as a template for subcloning to create yeast integrative plasmids (see below).

The sequence for Lon gene from *Mesoplasma florum* was based on the mf-Lon protein sequence obtained from NCBI database (GenBank: KM521209.1, Uniprot ID:Q6F160). A Su-9 mitochondrial targeting signal (amino acids 1-69 of the subunit 9 of the F0 ATPase of *Neurospora crassa*, including a mitochondrial processing peptidase site) as described in (*27*), and a 6xHis-3xFLAG tag were added to N- and C-termini, respectively.

A Protein Degradation Tag (PDT) (AANKNEENTNEVPTFMLNAGQANRRRV) was added to the C-terminus of GFP, and a Su-9 MTS at the N-terminus (*14*). Both constructs were codon optimized for expression in *Saccharomyces cerevisiae* (*S. cerevisiae*) and synthesized by the GeneART gene synthesis service at Thermo Fisher Scientific.

To construct yeast integrative plasmids, the MTS-mf-Lon and MTS-GFP-PDT constructs were first cloned in pVT100U-mtGFP between the *HindIII-NotI* restrictions sites, resulting in pVT100U-mtGFP-PDT (CD422) and pVT100U-mt-mf-Lon (CD423) plasmids. The pVT100U-mtGFP, pVT100U-mtGFP-PDT and pVT100U-mt-mf-Lon were then digested by *SphI* to release the *P*_*ADH1*_-*MTS-GFP-T*_*ADH1*_, *P*_*ADH1*_-*MTS-GFP-PDT*-*T*_*ADH1*_, and *P*_*ADH1*_-*MTS-MF-LON*-*T*_*ADH1*_ fragment, respectively. These were subsequently cloned into the *SphI* site of YIplac204 (GFP and GFP-PDT constructs), and YIplac128 (mf-Lon construct) integrative vectors (*28*).

The MTS was removed from YIplac204-mtGFP (CD424) and YIplac204-mtGFP-PDT (CD425) by amplifying the *P*_*ADH1*_ through Polymerase Chain Reaction (PCR) and the resulting DNA fragments were cloned between the *SalI-BglII* and *PciI-BglII* restriction sites of YIplac204-mtGFP and YIplac204-mtGFP-PDT, respectively, resulting in YIplac204-GFP (CD441) and YIplac204-GFP-PDT (CD442) plasmids.

For PDT-tagging of endogenous genes, *kanMX4* from pFA6-kanMX4 was cloned into the *NotI* site of YIplac204-mtGFP-PDT (CD425) plasmid. The resulting YIplac204-mtGFP-PDT-KanMX4 (CD436) plasmid was used as a template to amplify *PDT-kanMX4* using gene-specific primers for subsequent PDT tagging using established protocols for one step epitope tagging (*29*) (see Table S2).

### Strain

All strains used in this manuscript are derived from CB67 (*W303, MATa, ade2-1, trp1-1, can1-100, leu2-3*,*112, his3-11*,*15, ura3, GAL, psi+, RAD5*) and listed in Table S1. Standard transformation protocols were used. Each experiment was performed with at least two independent colonies.

To obtain mitochondrial GFP visualization strains, wild type yeast cells (CB67) were transformed with the *Bsu36I*-linearized YIplac204-mtGFP (CD424) or YIplac204-mtGFP-PDT (CD425) for integration at the *TRP1* locus. The resultant mito-GFP and mito-GFP-PDT strains were then transformed with *AflII*-linearized YIplac128-mt-mf-Lon (CD429) plasmid for integration at the *LEU2* locus. The same procedure was used to obtain strains for cytoplasmic GFP visualization except that cells were initially transformed with *Bsu36I*-linearized YIplac204-GFP (CD441) and YIplac204-GFP-PDT (CD442), i.e., constructs lacking MTS.

*MF-LON* was tagged C-terminally with mCherry using pFA6a-link-yomCherry-Kan as template as described in (*30*).

*PIM1* gene was knocked out in wild type or *P*_*ADH1*_-driven mito-mf-Lon strain by one step replacement of *PIM1* open reading frame (ORF) with *pim1Δ::natMX4*-cassette amplified from pCloneNat1 plasmid using primers flanking the *PIM1* ORF (Table S2). Strains containing the construct were selected on YPD agar plates supplemented with 100 mg/l of nourseothricin (Jena Bioscience, cat no. AB-101-10ML). The *pim1Δ*::mito-mf-Lon strains were created similarly except that an *MTS-MF-LON-LEU2* DNA fragment, amplified from the YIplac204-mt-mf-Lon plasmid, was used to replace *PIM1* ORF. The correct transformants were selected on solid minimal medium lacking leucine. All integrations were confirmed by PCR.

Pif1 and Apn1 were tagged with PDT at their C-terminus in wild type or mito-mf-Lon strains in two steps. First, the sequence encoding a 6xHA tag was introduced at the 3’ends of endogenous *PIF1* and *APN1* genes using *HIS3*-marked pYM15 plasmid as a template for the transformation construct (*6*). Second, the 6xHA tagged genes were PDT-tagged using the *PDT-kanMX4* cassette amplified from YIplac204-mtGFP-PDT-kanMX4 (CD436), replacing the *HIS3* marker with *kanMX4*. The correct strains were selected on YPD agar plates supplemented with 200 mg/l of G418 (Thermo Fisher Scientific, cat no 10131019), and confirmed by PCR.

### Media

Rich medium (Yeast Peptone Dextrose, YPD) consisted of 1% yeast extract (Thermo Fisher Scientific, product number 212750), 2% peptone (Thermo Fisher Scientific, product number 211677) and 50 mg/l adenine (Merck, product number A9126-100G), and was supplemented with appropriate carbon source [2% glucose (non-respiratory medium) or 3% glycerol (respiratory medium)], as indicated. Standard minimal medium (SMM) consisted of 0.67% yeast nitrogen base without amino acids (Merck, product number Y0626), supplemented with all standard amino acids [bought from Sigma (now Merck)] at 76 mg/l except leucine, which was added to 380 mg/l; adenine to 19 mg/l, and PABA to 7.6 mg/l, and 2% glucose was added as a carbon source (*31*).

For induction of PDT degradation, the composition of SMM was the same as described above, with the exception of adenine and glucose, which varied depending on the condition –”un-inducing” medium contained 50 mg/l adenine and 2% glucose, while “inducing” medium contained 2 mg/l adenine and 1% glucose.

For experiments when mitochondrial DNA (mtDNA) was visualized by Sybr Green I staining, SMM was supplemented with 340 mg/l isoleucine, 550 mg/l of leucine, and 430 mg/l of valine, to prevent parsing of nucleoids (*32*).

### Growth conditions

All experiments were initiated from fresh colonies obtained from glycerol stocks stored at − 80° C and grown on solid YPD media overnight at 30° C. Liquid cultures were grown at 30° C and 180 rotations per min (rpm). Each experiment was repeated at least twice.

For GFP visualization, fresh colonies were inoculated in 5 ml in SMM, and grown in 50 ml Falcon tubes for 16-18 hours, and samples were collected for GFP visualization at indicated timepoints (see Microscopy).

For growth-rate analysis, cells were inoculated in SMM at indicated optical densities (OD_600_), and growth was estimated by OD_600_ measurements at indicated timepoints.

### Induction of PDT degradation

For adenine titrations (Fig. 3, fig. S2B), mito-GFP + mito-mf-Lon-mCherry and mito-GFP-PDT + mito-mf-Lon-mCherry cells were grown to logarithmic growth (log phase, OD_600_ = 1.2-1.5), after which cells were pelleted, washed with 1x PBS buffer, and resuspended in SMM containing 1% glucose and 0 mg/l, 2 mg/l, or 20 mg/l adenine at a cell concentration of OD_600_ = ∼3, and thereafter grown for 3 hours at 30° C and 180 rpm. Samples were collected at indicated timepoints and imaged as described below (see Microscopy).

For induction of Apn1-PDT and Pif1-PDT degradation, cells were inoculated in 100 ml SMM supplemented with 50 mg/l adenine and 2% glucose at OD = 0.2 and grown for 6 hours to obtain a log phase culture (OD_600_ = 1). The cultures were then expanded by re-inoculation in 400 ml of fresh, un-inducing medium and grown overnight to an OD_600_ of 7-8. Cells were subsequently pelleted by centrifugation (4000 rcf, 5 min, room temperature), washed once with sterile water, and resuspended in 400 ml of inducing medium (SMM supplemented with 2 mg/l adenine and 1% glucose). To ensure that PDT degradation was induced, mito-GFP + mito-mf-Lon and mito-GFP-PDT + mito-mf-Lon cells were grown in parallel with the experimental strains, and PDT-degradation was analyzed by microscopy. Furthermore, the accumulation of red pigment resulting from adenine depletion served as a visual reference of diauxie (*20, 33*), which usually occurred 3-4 h after the shift to inducing medium.

For induction in wild type, *P*_*ADH1*_-driven mito-mf-Lon, and *pim1Δ*::mito-mf-Lon strains, cells were first grown in 20 ml un-inducing medium to reach log phase (OD_600_ = 1). Subsequently, cells were reinoculated in 50 ml un-inducing medium and grown overnight to an OD_600_ = 8-10 (30° C, 180 rpm, doubling time = 1.667 h). Finally, cells were washed and shifted to 50 ml of inducing medium and grown for ∼4 h. Samples were collected before and after induction for estimation of mtDNA by quantitative real time PCR (qPCR).

### Visualization of mtDNA

Wild type, *P*_*ADH1*_-driven mito-mf-Lon, *pim1Δ*::mito-mf-Lon strains were grown overnight in SMM with excess isoleucine, leucine and valine (SMM+ILV) to OD_600_ = 8-10 (see Media). The excess branched chain amino acids did not affect mf-Lon-induced PDT degradation (data not shown). One milliliter of overnight culture was collected and washed with PBS using centrifugation (3000 rpm, 2 min), and incubated for 30 s at room temperature in 500 µl staining solution containing 2.5 µl of Sybr Green I (Thermo Fisher Scientific, S7563) in PBS. Finally, cells were washed twice with PBS using centrifugation (3000 rpm, 2 min) before imaging in SMM+ILV medium.

### Microscopy

70 µl of cells were immobilized on glass bottom wells of imaging plates (Mobitec, Imaging Plate 96 CG 1.5, Order #: 5242-20) coated with Concanavalin A (0.45 mg/ml, Sigma) and overlaid with appropriate media. For confocal microscopy (Figs. 1B, 2B and E, figs. S1A and S4D), cells were imaged in a full-size temperature-controlled incubator (set at 30° C throughout the experiment) on a Nikon Ti Eclipse microscope equipped with a spinning disk confocal (Yokagawa CSU-X1), an ALC laser controller, and an Andor DU-897 X-3655 EM-CCD camera (Pixel size 16 µm. QE 95%), using a Nikon 100x/1.4 PlanApo oil objective. A 1.2x lens was inserted in the emission light path to fulfil the Nyquist sampling theorem with the camera used. The acquisition settings for GFP were 488 nm diode laser, 9.4%, 50 ms exposure time, 7.5 µm Z-stack with 0.3 µm interval, and for mtDNA staining with Sybr Green I (fig. S3A), 488 nm diode laser at 9.8% power, 20 ms exposure time.

For widefield microscopy (Figs. 1D, 2A, 3A-C, figs. S2A and S4C), cells were imaged in the same environmental conditions and on the same microscope as above, but using an Andor Zyla 4.2+ sCMOS camera (pixel size 6.45 µm, QE 82%) without the 1.2x lens,, a Lumencore SpectraX LED light source, a Suttter external emission filter wheel, a 60x/1.2 PlanApo water objective, and a 1.5x magnification lens inserted, in order to fulfill Nyquist sampling theorem. The acquisition settings were the following: Lon-mCherry – excitation filter 555 nm at 84%, emission filter 630/90 nm, exposure time 400 ms; mito-GFP – excitation filter 470 nm at 22%, emission filter of 525/30 nm, exposure time 30ms; cyto-GFP – excitation filter 470 nm at 50%, emission filter 525/30 nm, exposure time 200 ms, 5.55 µm Z-stack with 0.37 µm interval.

### Image analysis

Images of mitochondrial GFP captured by widefield microscopy were deconvolved using the FAST deconvolution function of NIS elements software (Nikon Instruments, Tokyo, Japan). All images were colorized in the Fiji software and shown as maximum intensity Z-projections. The brightness and contrast values were set equally along all samples.

For quantification of fluorescent signals, yeast cell segmentation was performed by YeastSpotter (http://yeastspotter.csb.utoronto.ca) (*34*). Cell quantification was performed in Fiji (*35*) on maximum intensity Z-projections. Cells that had area less than 10 µm^2^ or circularity less than 0.85 were excluded from analysis. For mito-GFP/GFP-PDT and cyto-GFP/GFP-PDT expressing cells, intensity was estimated using integrated density in the whole cell after background subtraction. To minimize experiment-to-experiment variability, the values were normalized to the median level displayed by cells expressing GFP only.

For nucleoid analysis after Syber Green I staining, the spots segmentation was performed in Fiji on maximum intensity Z-projections after computing Laplacian image using FeatureJ plugin (http://imagescience.org/meijering/software/featurej/). To exclude noise/debris and occasional weakly stained nucleus DNA, spots that displayed area less than 0.06 µm^2^ or median integrated density less than 500 a.u (arbitrary units). were excluded from analysis. To adjust nucleoid number in occasional aggregates, we divided each spot integrated density by the median spot integrated density found on the image and rounded the result to the nearest integer above zero. To minimize experiment-to-experiment variability, values were normalized to the average level displayed by the wild-type cells (CB67). Statistical analysis was performed by *Hierarch* resampling Python-based module (*36*), and a *P* value of less than 0.05 was considered statistically significant. Graphical data presentation was done by GraphPad Prism 9.0 (www.graphpad.com).

### Estimation of mtDNA copy number by qPCR

Wild type, *P*_*ADH1*_-driven mito-mf-Lon, *pim1Δ*::mito-mf-Lon strains were induced for PDT degradation, and 2 ml of culture was harvested before and after shifting cells to the inducing media. Total DNA was isolated by phenol:chloroform:isoamyl alcohol and cold ethanol precipitation method. Relative quantity of mtDNA was estimated by qPCR using primers against *COX2* and *ACT1* (Table S2) and a standard curve generated from total DNA isolated from an uninduced culture of wild type with Fast SYBR Green (Applied Biosystems, 4385612) according to manufacturer’s protocol. Unpaired, 2-tailed t-test was performed to analyze data with error bars representing standard deviations from 3 independent experiments.

### Isolation of mitochondria

Mitochondria were prepared from yeast cells by differential centrifugation method as described in (*37*). Briefly, 400 ml of Apn1-PDT and Pif1-PDT cells, expressing mf-Lon from the *ADH1* promoter, or from *PIM1* locus, induced for PDT degradation, were harvested (5000 rpm, 10 min, room temperature), resuspended in 2 ml of MP1 buffer for every gram of cell pellet (100 mM Tris, 10 mM DTT), and incubated (30° C, 10 min). The cells were subsequently washed with 1.2 M sorbitol and resuspended in MP2 [1.2 M Sorbitol, 0.02 M potassium phosphate buffer, pH 7.4, 20T Zymolyase (Seikagaku Biobusiness, product code 120491) at 3 mg/g cell pellet] using 6.7 ml per gram of cell pellet, followed by incubation (30° C, 60 min) to create sphaeroplasts which lack the yeast cell wall. At this point, a 50 µl cell extract sample was taken and boiled for 5 min with 50 µl of 5x SDS loading buffer, and stored at −20° C until analysis by Western Blotting.

The remaining sphaeroplasts were centrifuged (4500 rpm, 6 min at 4° C), and resuspended in MP3 buffer (10 mM Tris pH 7.4, 1 mM EDTA, 1 mM PMSF, 0.6 M sorbitol) using 13.4 ml per gram of cell pellet. This sphaeroplast suspension was then homogenized using a tissue grinder (Fisherbrand, product code 10331592), centrifuged (4500 rpm, 6 min at 4° C), and the supernatant was cleared of any remaining cell debris by two additional centrifugations (4500 rpm, 6 min, at 4° C). The resulting supernatant was re-centrifuged (20000 g, 25 min at 4° C), and the pellet, representing a crude preparation of mitochondria was resuspended in SH buffer (0.6 M sorbitol, 20 mM HEPES pH 7.4) to a concentration of 10 mg/ml. These were finally snap frozen in liquid nitrogen and stored at −80° C.

### Proteinase K treatment of mitochondria

600 µg of mitochondria were harvested in two 1.5 ml Eppendorf tubes (300 µg each) by spinning (10000 g, 10 min at 4° C) in a tabletop centrifuge. 40 µl of SH buffer was added to the pellet in each tube and incubated on ice for 10 min. Then, 5 µl of 0.5 mg/ml of Proteinase K (Sigma RPROTK-RO, dissolved in SH buffer at 50 µg/ml) was added to one of the tubes, and both tubes were incubated on ice for another 10 min. Thereafter, 5 µl of 100 mM PMSF was added to both tubes, that were subsequently centrifuged (21000 g, 10 min, at 4° C). The resulting mitochondrial pellets were washed twice in SHKCL buffer (0.6 M sorbitol, 20 mM HEPES pH 7.4, 150 mM KCL) by centrifugation (21000 g, 10 min at 4° C), and finally resuspended in 100 µl of 2.5x SDS loading dye. These were then boiled for 10 min, supernatant collected by centrifugation (14000 rpm, 10 min, 4° C) and stored at −20° C until analysis by Western Blotting.

### Western blotting

For cell extract samples from induced culture, 50 µl sphaeroplasts suspension was boiled directly with 2.5x SDS loading buffer before analysis. The cell extract protein sample from uninduced culture was prepared by standard trichloroacetic acid precipitation method. Briefly, 250 µl of 20% TCA was added to 10 ml of pelleted cell culture, and the mixture was vortexed in the presence of glass beads to disrupt the cells. The suspension was pelleted at 13000 rpm, 4 min at room temperature, and washed twice with ice-cold 100% ethanol and air-dried. The dried pellets were subsequently resuspended in 2.5x SDS loading buffer and boiled for 5 min. Cell debris were removed by centrifugation (14000 rpm, 10 min, 4° C) and the supernatant was stored at −20° C before analysis by Western Blotting according to standard protocols. The following antibodies were used: PDT-tagged proteins - anti-HA antibody (Sigma, cat# 11666606001, dilution 1:1000), mf-Lon - anti-Flag (Sigma-Aldrich, cat# F1804-5 MG, dilution 1:3000), histone - anti-H3 (Abcam, cat# ab1791, dilution 1:1000), Cox2 – anti-MTCO2 (Abcam, cat# ab110271, dilution 1:1000); the following antibodies were gifts from Martin Ott: Mrpl4 – anti-Mrpl4 (dilution 1:2000), Mrpl36 – anti-Mrpl36 (dilution 1:500), Mrpl40 – anti-Mrpl40 (dilution 1:2000), and Mdh1 – anti-Mdh1 (dilution 1:10000).

## Supplementary Text

### Choice of the PDT tag

Most bacterial species employ more than one protease to degrade the ssrA tag, and owing to the evolutionary conservation, proteases display inter-species promiscuity in tag detection, making controllable protein degradation challenging (*13, 14, 38*–*40*).

Since the *Escherichia coli* (*E. coli*) Lon can degrade mf-Lon ssrA tag, and can functionally complement the yeast Lon, we aimed to use a variant of mf-Lon ssrA tag that is unidentifiable by the *E. Coli* -Lon [pdt#3 (AANKNEENTNEVPTFMLNAGQANRRRV)], to avoid interference by yeast endogenous Lon (*13, 14, 41*). Throughout the manuscript, we referred to the pdt#3 tag as PDT.

### Optimization of PDT degradation

We considered three parameters to mimic diauxic shift and PDT degradation; high cell concentration, glucose concentration and adenine concentration.

In synthetic minimal medium which contains 18.94 mg/l adenine, a W303 strain which is an adenine auxotroph starts decelerating cell growth upon reaching an OD_600_ ∼4-5. Therefore, cells were usually resuspended in the inducing medium at a cell concentration of OD higher than 4.

Cells were grown in 1% glucose to shift cellular metabolism towards respiration, which occurs during diauxic shift (*15*).

Finally, we chose 2 mg/l of adenine for PDT degradation since this concentration allowed cell division, as judged by budding index, in contrast to complete deprivation of adenine (0 mg/l) which, although inducing PDT degradation faster, impaired cellular growth.

---

**Fig S1.**
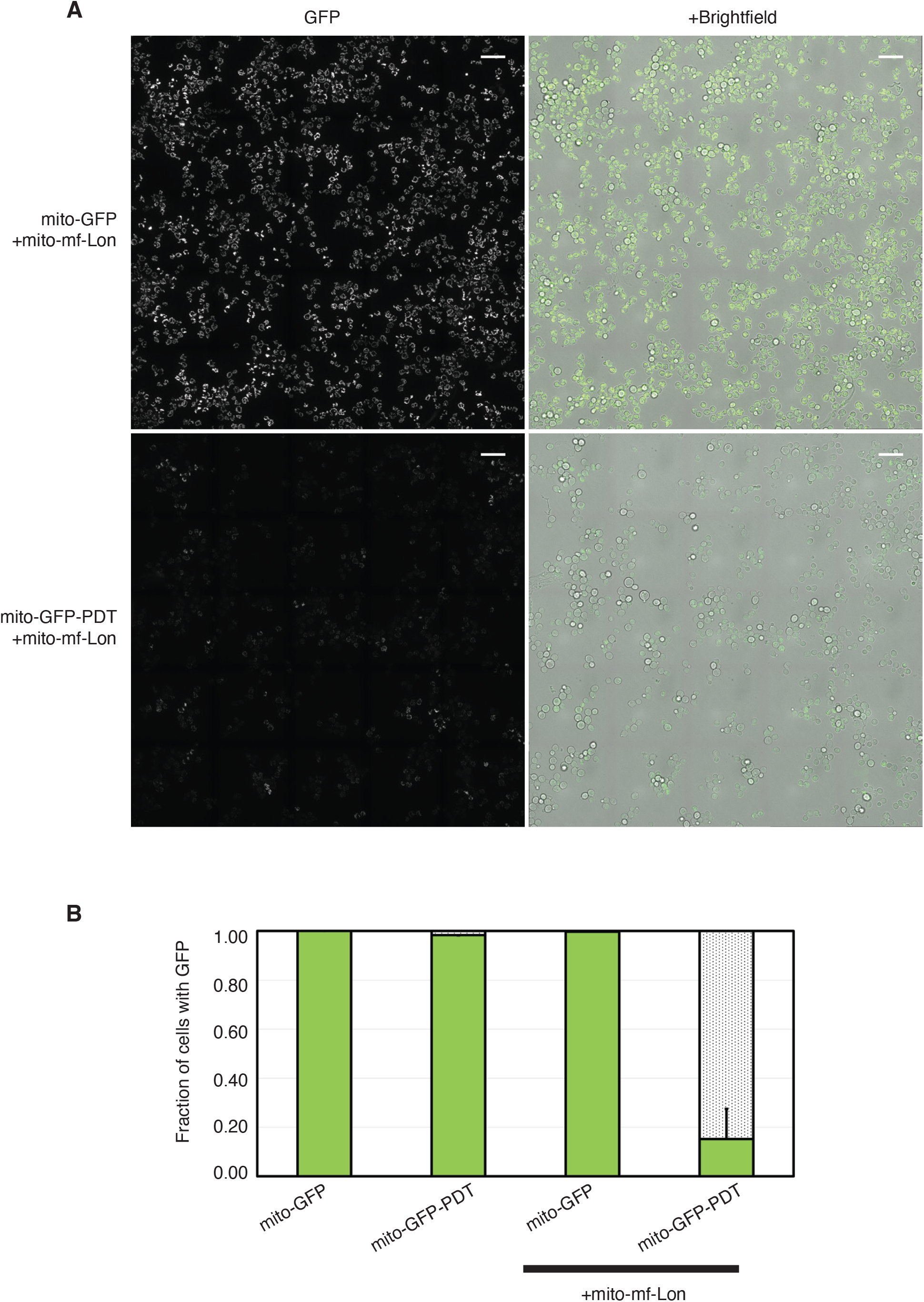
mf-Lon induces mitochondrial GFP-PDT degradation. **(A)** Large field image of the GFP signal in mito-GFP mito-mf-Lon, and mito-GFP-PDT mito-mf-Lon cells. Scale bar = 20 μm. Cells were grown in minimal dextrose medium and imaged when they reached deceleration growth phase. **(B)** Fraction of cells with mitochondrial GFP in the indicated strains. Error bars represent standard deviation from 3 independent colonies. Cells were grown in minimal medium and examined after 16-18h of inoculation (overnight culture).

**Fig S2.**
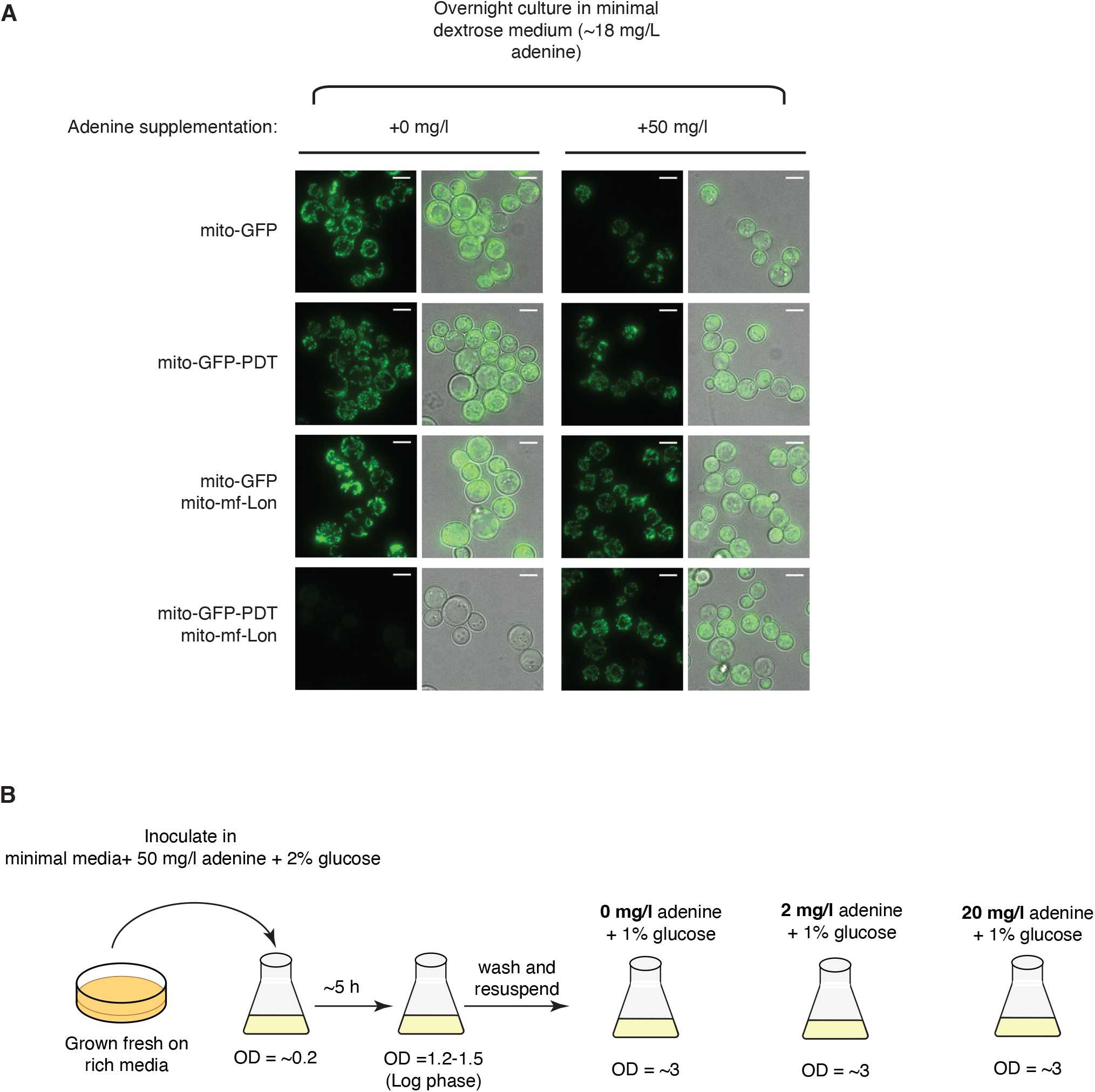
Adenine supplementation to minimal medium prevents mitochondrial GFP-PDT degradation by mf-Lon. **(A)** Indicated strains growing in standard minimal dextrose medium (comprising ∼18 mg/l adenine) were left untreated (left panel), or supplemented with adenine (right panel). Representative live cell images of an overnight culture are shown. Scale bar = 5 µm. **(B)** Schematic for adenine titration experiment in Figure 3. Cells expressing mitochondrially targeted GFP and mf-Lon-mCherry, or GFP-PDT and mf-Lon-mCherry growing in 50 mg/l adenine and 2% glucose (un-inducing media) were harvested, washed and resuspeneded in minimal medium supplemented with 1% glucose and 0, 2, or 20 mg/l adenine. Samples were taken every hour and examined for degradation of GFP-PDT by widefield microscopy.

**Fig S3.**
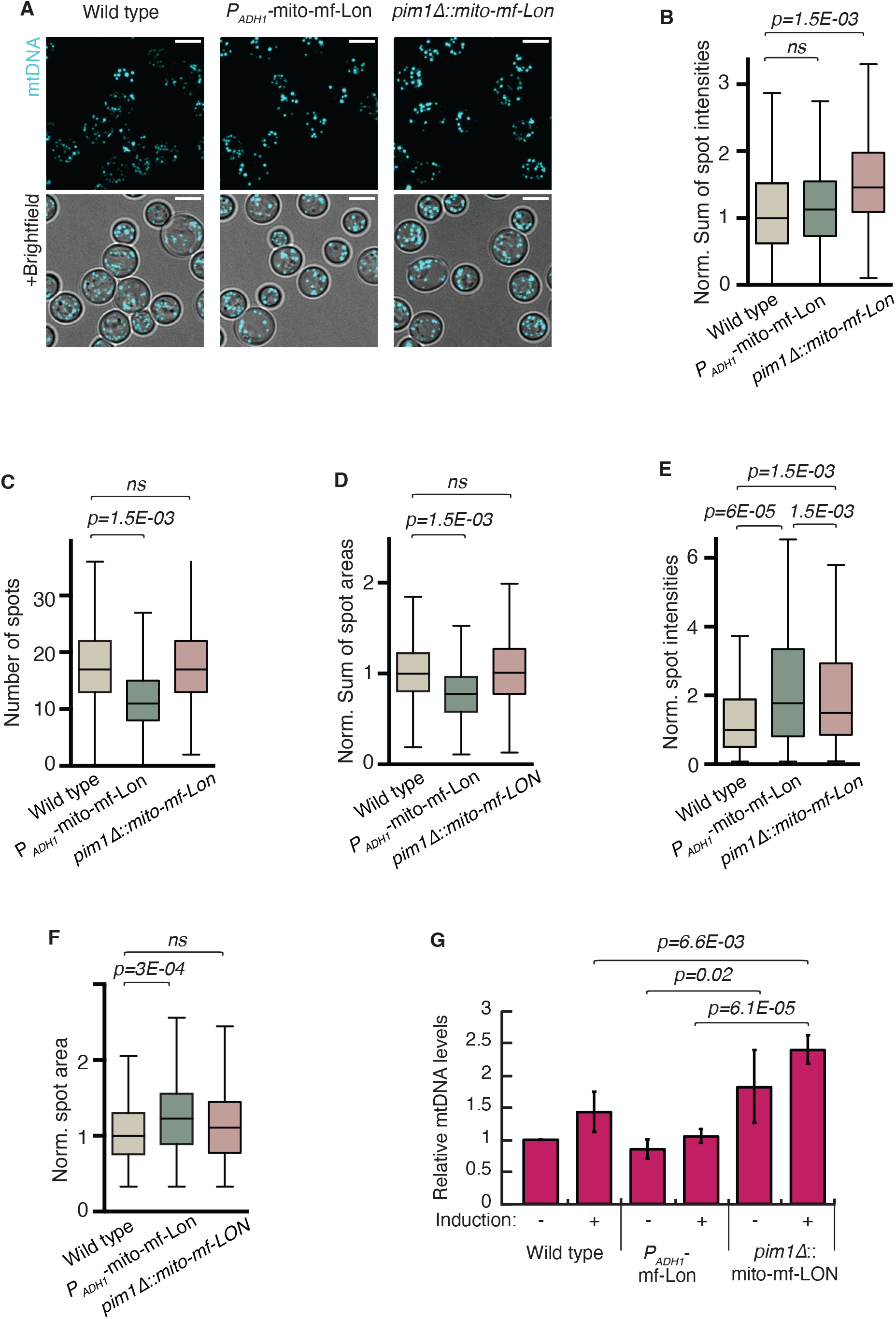
Visualization of mitochondrial nucleoids in wild type, *P*_*ADH1*_-driven mf-Lon, or *pim1Δ*::mito-mf-Lon cells. (A) Representative images of Sybr Green I-stained mtDNA indicated strains grown overnight in standard minimal medium. **(B-F)** Quantification of mtDNA spots detected by Sybr Green I. Sum of spot intensities in the cell, representing total mitochondrial DNA content per cell **(B)**, number of spots, representing number of nucleoids per cell **(C)**, sum of spot areas **(D)**, spot intensities **(E)**, and area of individual spots **(F)** were compared by hierarchical resampling. Boxes show interquartile range and whiskers at 10% and 90% range. Values in (B), (D), (E) and (F) were normalized to the wild type median. *P*-values (*p*) more than 0.05 were considered non-significant (*ns*). (G) WT, *P*_*ADH1*_-driven mito-mf-Lon, and *pim1Δ*::mito-mf-Lon cells were induced for PDT degradation, and the levels of mtDNA relative to uninduced sample of WT were determined by quantitative real time PCR. An unpaired, two-tailed students t-test was performed to determine the confidence interval. *P*-values (*p*) are indicated.

**Fig S4.**
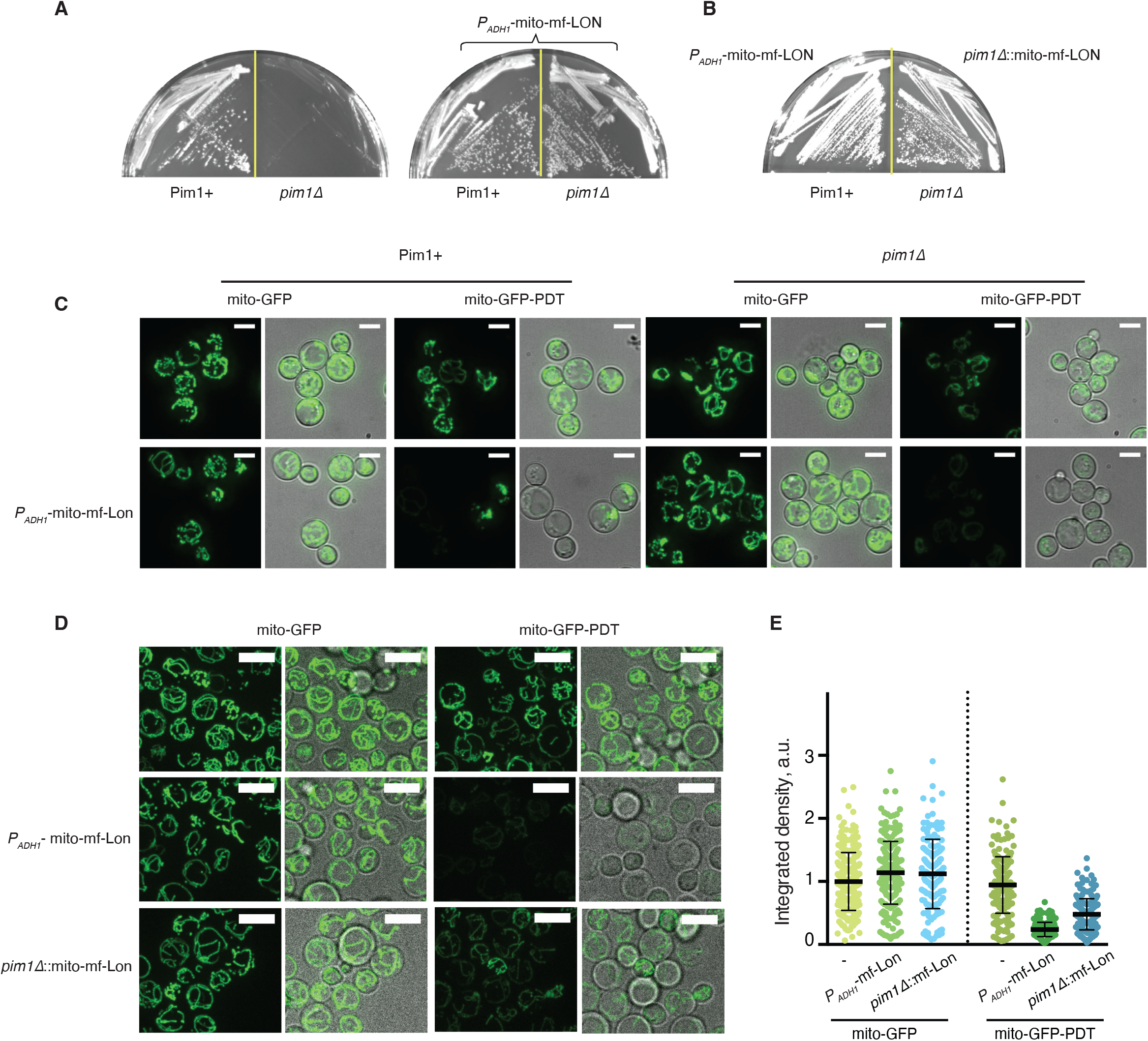
Lon-induced PDT degradation occurs independently of yeast endogenous Lon, Pim1. **(A-B)** mf-Lon rescues respiratory defect of *pim1Δ* cells. **(A)** Strains with or without wild type *PIM1*, either independently or co-expressing *P*_*ADH1*_-driven mito-mf-Lon, were grown on solid respiratory medium, and photographed after incubation at 30°C for 4 days. **(B)** Same as (A), but with strains that expressed mf-Lon from the *ADH1* promoter, or from the *PIM1* locus. **(C-D)** PDT degradation is specific to mf-Lon. **(C)** Cells expressing mito-GFP or mito-GFP-PDT, either independently or co-expressing *P*_*ADH1*_-driven mito-mf-Lon, in the presence or absence of wild type *PIM1* were grown overnight in standard minimal medium and examined for GFP-PDT degradation. Representative live cell images are shown. **(D)** Cells expressing mito-GFP or mito-GFP-PDT, either independently or co-expressing *P*_*ADH1*_-controlled mito-mf-Lon, or *pim1Δ*::mito-mf-Lon were examined for PDT degradation similarly to (C). Note that both mito-GFP and mito-GFP-PDT are also expressed from the *ADH1* promoter. Scale bar=5µm. **(E)** Quantification of GFP signal in the cells shown in (D), expressed as arbitrary units (a.u.). Each dot represents an individual cell. Data depict median and interquartile range and are normalized to the median intensity in mito-GFP strain.

**Fig. S5.**
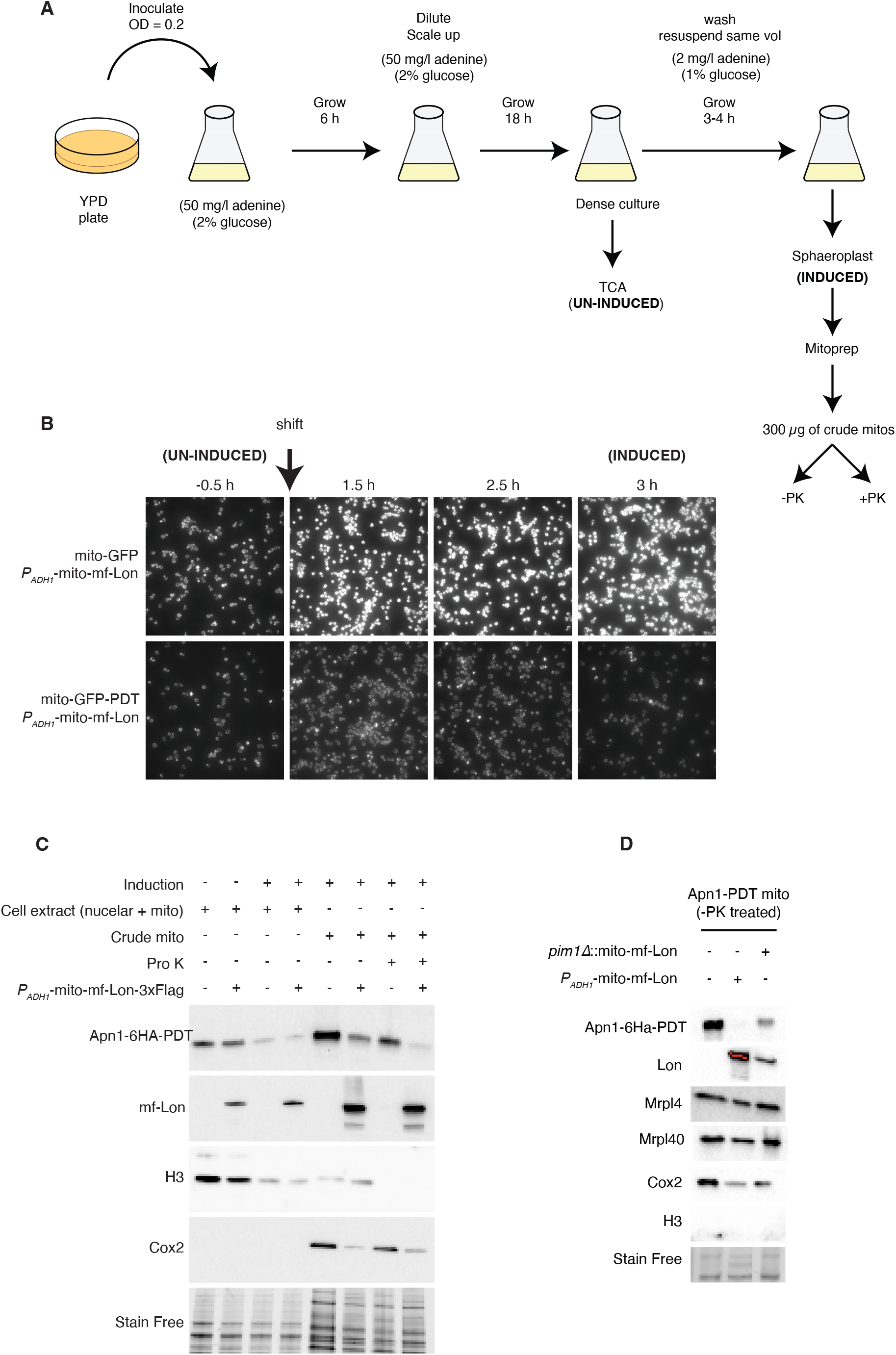
mf-Lon protease degrades yeast endogenous mitochondrial proteins tagged with PDT. **(A)** Schematic of induction experiment (also see Materials and Methods for details). **(B)** Representative images of mito-GFP plus mito-mf-Lon and mito-GFP-PDT plus mito-mf-Lon cells imaged 30 mins before (un-induced) and up to 3 hours after the shift in inducing medium (induced), grown in parallel with the experimental strains, to ensure that the media induced PDT degradation. **(C)** Cells expressing Apn1-PDT with or without *P*_*ADH1*_-controlled mf-Lon were induced for PDT-degradation as shown in (A). Cell extracts from uninduced and induced cultures, and mitochondria from induced samples, were analyzed by Western Blotting for indicated proteins. Proteinase K treatment was performed on mitochondria to remove nuclear contamination. **(D)** Mitochondrial samples as shown in Figure 4B were analysed by Western Blotting for additional mitochondrial and nuclear proteins. Absence of H3 indicates a cleaner mitochondrial preparation. Stain free image serves as a loading control. The sizes of the proteins are: Apn1-PDT (anti-HA), ∼60 kDa; mf-Lon (anti-Flag), ∼100 kDa; H3 (anti-H3), ∼15 kDa; Cox2 (anti-MTCO2), ∼35 kDa, Mrpl4 (anti-Mrpl4), ∼44 kDa; Mdh1 (anti-Mdh1), 35.6 kDa; and Mrpl40 (anti-Mrpl40), ∼40 kDa.

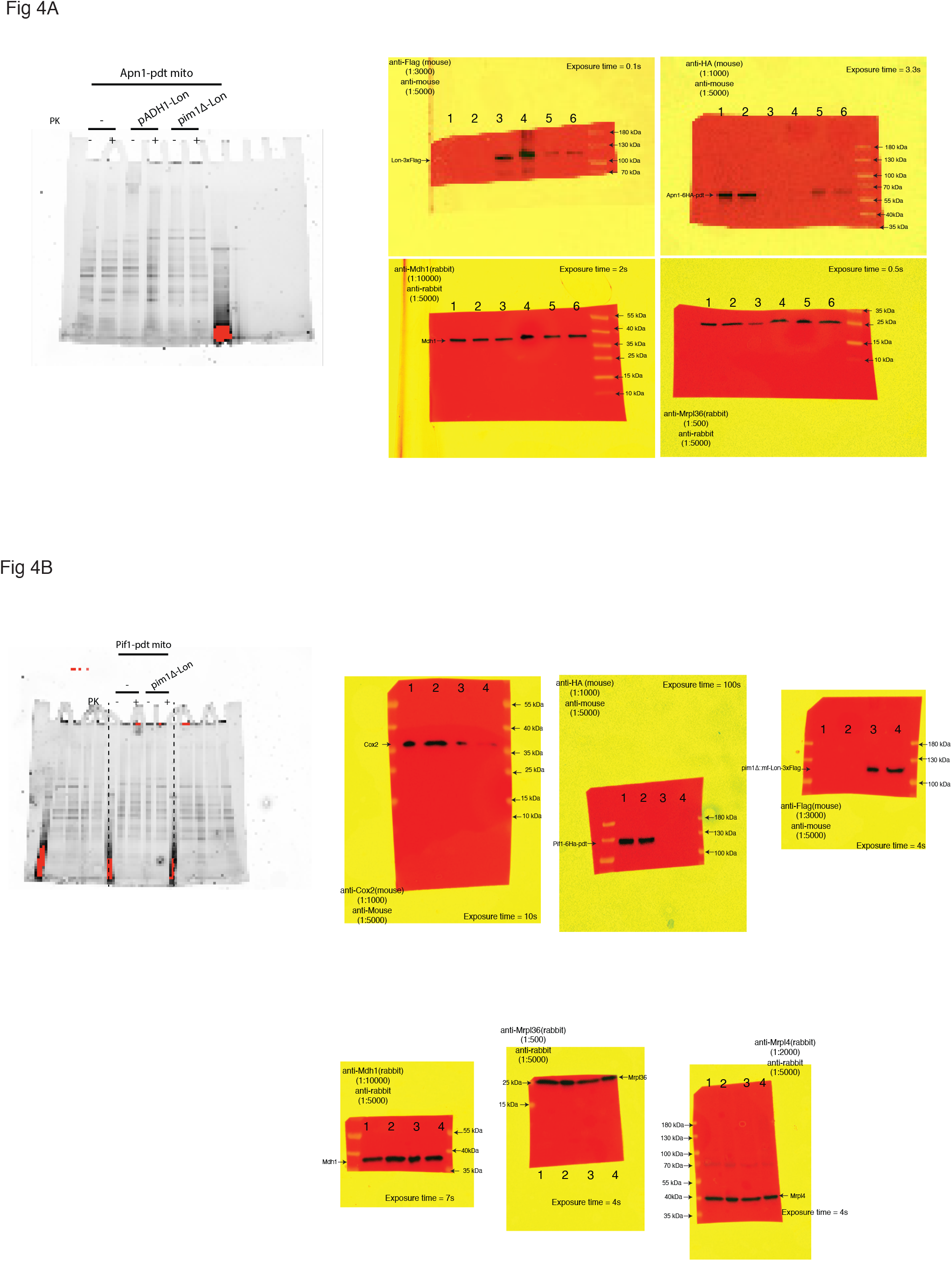
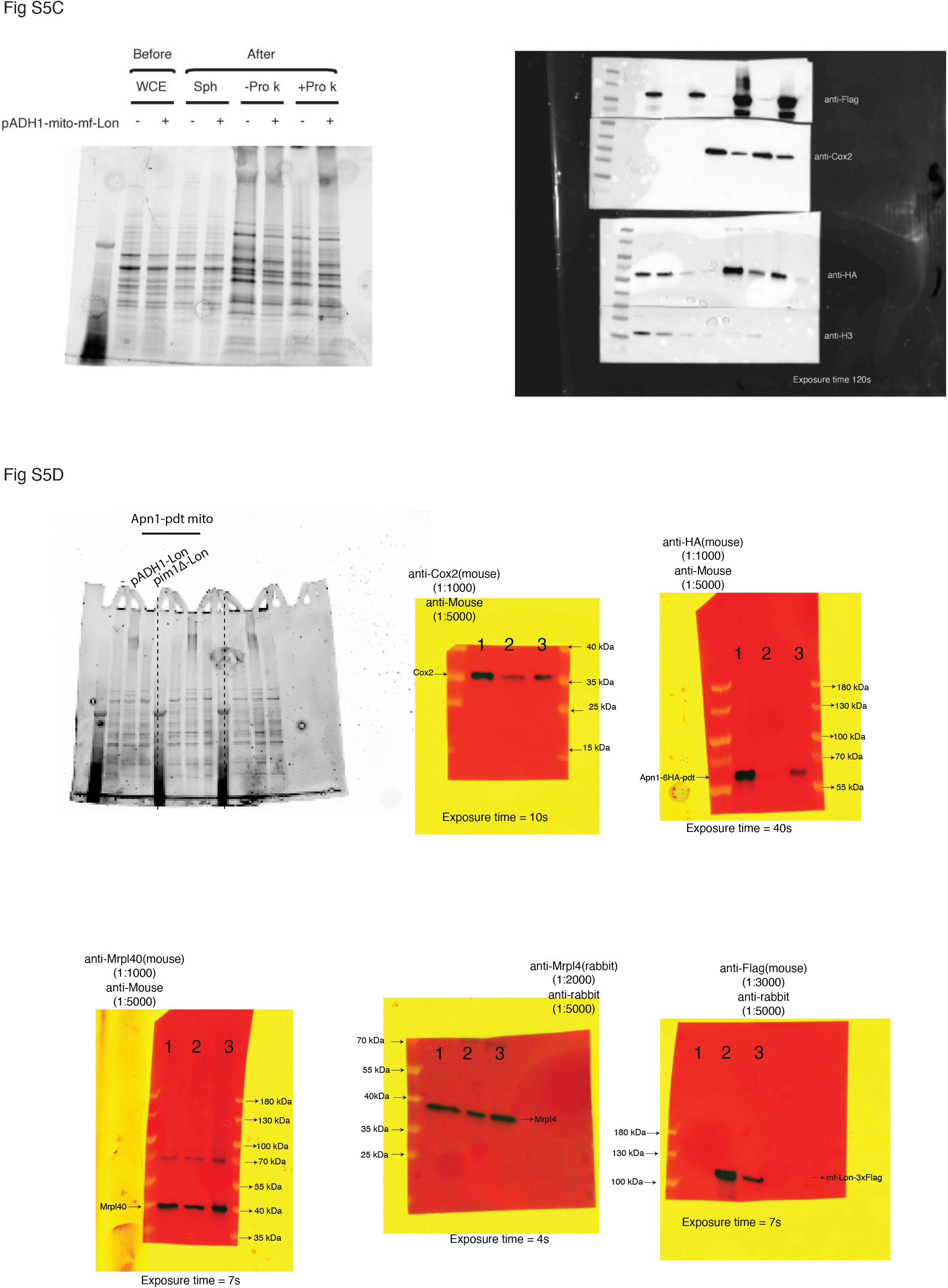

**Table S1.** All strains listed were derived from *RAD5* W303 strain, CB67 (*MATa, ade2-1, trp1-1, can1-100, leu2-3*,*112, his3-11*,*15, ura3, GAL, psi+, RAD5*), denoted wild type in the manuscript.

| Name | Identifier | Genotype |
| --- | --- | --- |
| mito-GFP | CB3699 | *trp1-1::TRP1-P<sub>ADHI</sub>-MTS (SU9<sup>1-69</sup>)-GFP-T<sub>ADHI</sub>*<br>(*Yiplac204*) |
| mito-GFP-PDT | CB3701 | *trp1-1::TRP1-P<sub>ADHI</sub>-MTS (SU9<sup>1-69</sup>)-GFP-PDT#3-T<sub>ADHI</sub>*<br>(*Yiplac204*) |
| mito-GFP<br>mito-mf-Lon | CB3713 | *trp1-1::TRP1-P<sub>ADHI</sub>-MTS (SU9<sup>1-69</sup>)-GFP-T<sub>ADHI</sub>*<br>(*Yiplac204*), *leu2-3::LEU2-P<sub>ADHI</sub>-MTS (SU9<sup>1-69</sup>)-MF-LON-6xHIS-3xFLAG-T<sub>ADHI</sub>*<br>(*Yiplac128*) |
| mito-GFP-PDT<br>mito-mf-Lon | CB3714 | *trp1-1::TRP1-P<sub>ADHI</sub>-MTS (SU9<sup>1-69</sup>)-GFP-PDT#3-T<sub>ADHI</sub>*<br>(*Yiplac204*), *leu2-3::LEU2-P<sub>ADHI</sub>-MTS (SU9<sup>1-69</sup>)-MF-LON-6xHIS-3xFLAG-T<sub>ADHI</sub>*<br>(*Yiplac128*) |
| cyto-GFP | CB3767 | *trp1-1::TRP1-P<sub>ADHI</sub>-GFP-T<sub>ADHI</sub>* (*Yiplac204*) |
| cyto-GFP-PDT | CB3769 | *trp1-1::TRP1-P<sub>ADHI</sub>-GFP-PDT#3-T<sub>ADHI</sub>* (*Yiplac204*) |
| cyto-GFP<br>mito-mf-Lon | CB3782 | *trp1-1::TRP1-P<sub>ADHI</sub>-GFP-T<sub>ADHI</sub>* (*Yiplac204*),<br>*leu2-3::LEU2-P<sub>ADHI</sub>-MTS (SU9<sup>1-69</sup>)-MF-LON-6xHIS-3xFLAG-T<sub>ADHI</sub>*<br>(*Yiplac128*) |
| cyto-GFP-PDT<br>mito-mf-Lon | CB3783 | *trp1-1::TRP1-P<sub>ADHI</sub>-GFP-PDT#3-T<sub>ADHI</sub>* (*Yiplac204*),<br>*leu2-3::LEU2-P<sub>ADHI</sub>-MTS (SU9<sup>1-69</sup>)-MF-LON-6xHIS-3xFLAG-T<sub>ADHI</sub>*<br>(*Yiplac128*) |
| mito-GFP<br>mito-mf-Lon-mCherry | CB3726 | *trp1-1::TRP1-P<sub>ADHI</sub>-MTS (SU9<sup>1-69</sup>)-GFP-PDT#3-T<sub>ADHI</sub>*<br>(*Yiplac204*),<br>*leu2-3::LEU2-P<sub>ADHI</sub>-MTS (SU9<sup>1-69</sup>)-MF-LON-mCherry-T<sub>ADHI</sub>-kanMx4*<br>(*pFA6a-link-yomCherry-Kan*) |
| mito-GFP-PDT<br>mito-mf-Lon-mCherry | CB3727 | *trp1-1::TRP1-T<sub>ADHI</sub>-MTS (SU9<sup>1-69</sup>)-GFP-P<sub>ADHI</sub>*<br>(*Yiplac204*), *leu2-3::LEU2-P<sub>ADHI</sub>-MTS(SU9<sup>1-69</sup>)-MF-LON-mCherry-T<sub>ADHI</sub>-kanMx4*<br>(*pFA6a-link-yomCherry-Kan*) |
| *pim1Δ*<br>mito-GFP | CB3786 | *pim1Δ::natMX4* (pCloneNat1), *trp1::TRP1-P<sub>ADH1</sub>-MTS* (*SU9<sup>1-69</sup>*)-GFP-*T<sub>ADH1</sub>* (*Yiplac204*) |
| *pim1Δ*<br>mito-GFP-PDT | CB3787 | *pim1Δ::natMX4* (pCloneNat1), *trp1-1::TRP1-P<sub>ADH1</sub>-MTS* (*SU9<sup>1-69</sup>*)-GFP-PDT#3-*T<sub>ADH1</sub>* (*Yiplac204*) |
| *pim1Δ*<br>mito-GFP<br>mito-mf-Lon | CB3790 | *pim1Δ::natMX4* (pCloneNat1), *trp1-1::TRP1-P<sub>ADH1</sub>-MTS* (*SU9<sup>1-69</sup>*)-GFP-*T<sub>ADH1</sub>* (*Yiplac204*), *leu2-3::LEU2-P<sub>ADH1</sub>-MTS* (*SU9<sup>1-69</sup>*)- *MF-LON* -6xHIS-3xFLAG-*T<sub>ADH1</sub>* (*Yiplac128*) |
| *pim1Δ*<br>mito-GFP-PDT<br>mito-mf-Lon | CB3791 | *pim1Δ::natMX4* (pCloneNat1), *trp1-1::TRP1-P<sub>ADH1</sub>-MTS* (*SU9<sup>1-69</sup>*)-GFP-PDT#3-*T<sub>ADH1</sub>* (*Yiplac204*), *leu2-3::LEU2-P<sub>ADH1</sub>-MTS* (*SU9<sup>1-69</sup>*)- *MF-LON* -6xHIS-3xFLAG-*T<sub>ADH1</sub>* (*Yiplac128*) |
| *pim1Δ::mito-mf-Lon*<br>mito-GFP | CB3921 | *trp1-1::TRP1-P<sub>ADH1</sub>-MTS*(*SU9<sup>1-69</sup>*)-GFP-*T<sub>ADH1</sub>* (*Yiplac204*), *pim1Δ::MTS*(*SU9<sup>1-69</sup>*)- *MF-LON* -6xHIS-3xFLAG-*LEU2* |
| *pim1Δ::mito-mf-Lon*<br>mito-GFP-PDT | CB3922 | *trp1-1::TRP1-P<sub>ADH1</sub>- MTS* (*SU9<sup>1-69</sup>*)-GFP-PDT#3-*T<sub>ADH1</sub>* (*Yiplac204*), *pim1Δ::MTS* (*SU9<sup>1-69</sup>*)- *MF-LON* -6xHIS-3xFLAG -*LEU2* |
| *pim1Δ::mito-mf-Lon* | CB3923 | *pim1Δ::MTS* (*SU9<sup>1-69</sup>*)- *MF-LON* -6xHIS-3xFLAG-*LEU2* |
| Apn1-PDT | CB3876 | *APN1-6xHA-PDT#3-kanMX4* |
| Apn1-PDT<br>*P<sub>ADH1</sub>*-mito-mf-Lon | CB3877 | *APN1-6xHA-PDT#3-kanMX4*, *leu2-3::LEU2-P<sub>ADH1</sub>-MTS* (*SU9<sup>1-69</sup>*)- *MF-LON* -6xHIS-3xFLAG-*T<sub>ADH1</sub>* (*Yiplac128*), |
| Apn1-PDT<br>*pim1Δ::mito-mf-Lon* | CB3954 | *APN1-6HA-PDT#3-kanMX4*, *pim1Δ:: MTS* (*SU9<sup>1-69</sup>*)-*MF-LON* N-6xHIS-3xFLAG-*LEU2* |
| Pif1-PDT | CB3871 | *PIF1-6xHA-PDT#3-kanMX4* |
| Pif1-PDT<br>*pim1Δ::mito-mf-Lon* | CB3955 | *PIF1-6HA-PDT#3-kanMX4*, *pim1Δ::MTS* (*SU9<sup>1-69</sup>*)-*MF-LON* -6xHIS-3xFLAG-*LEU2* |
| *P<sub>ADH1</sub>*-mito-mf-Lon | CB3730 | *leu2-3::LEU2-P<sub>ADH1</sub>-MTS (SU9<sup>1-69</sup>)- MF-LON -*<br>*6xHIS-3xFLAG -T<sub>ADH1</sub> (Ylplac128)* |

**Table S2.**
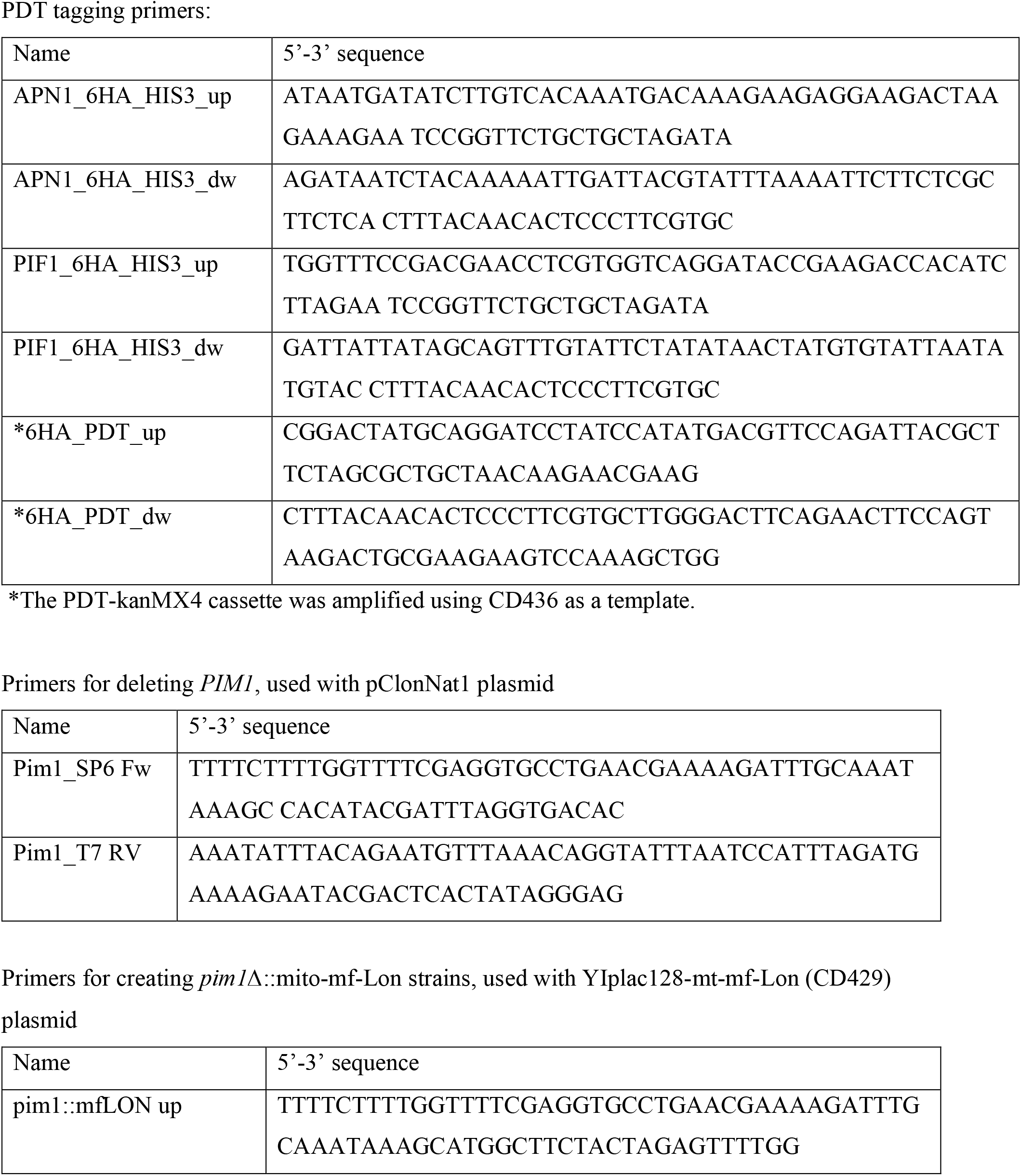
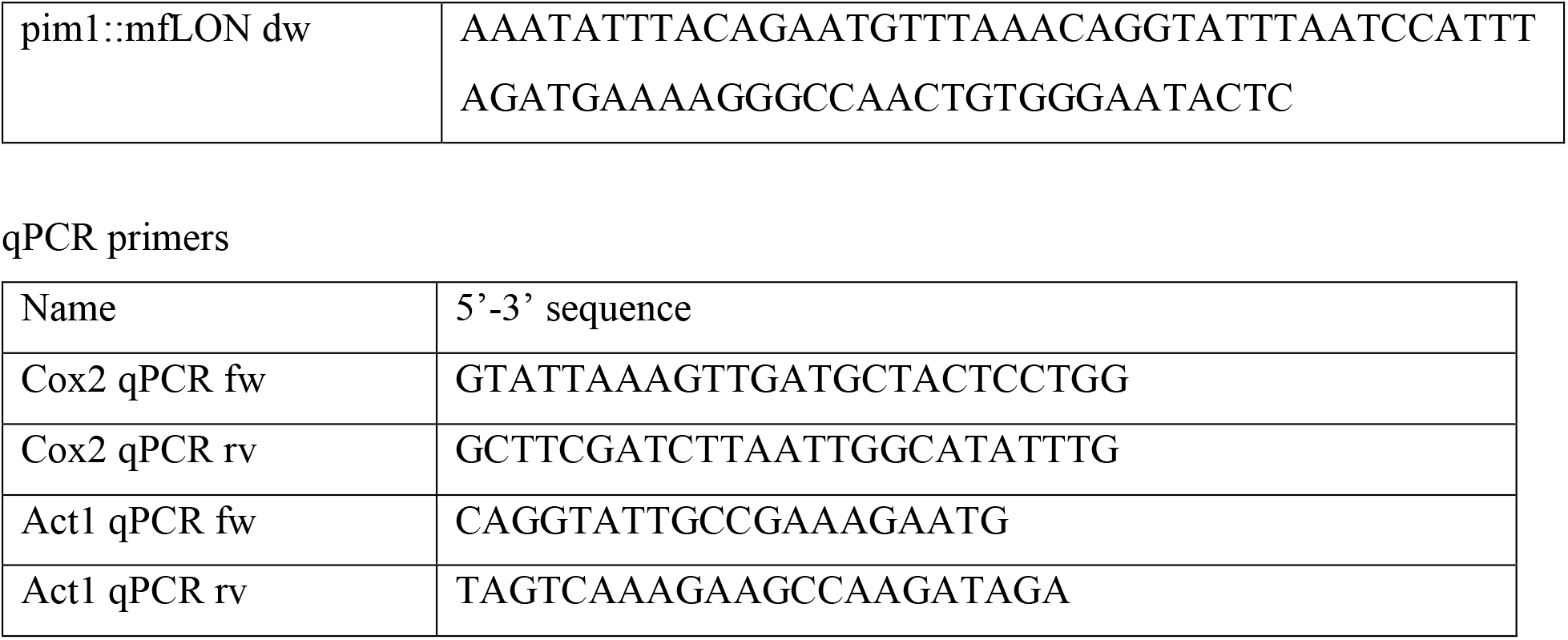
Primers used in the study.

## References and notes (including Supplementary Materials)

1. M. W. Gray, Mitochondrial Evolution. Csh Perspect Biol. 4, a011403 (2012).

2. J. Nunnari, A. Suomalainen, Mitochondria: In Sickness and in Health. Cell. 148, 1145–1159 (2012).

3. A. Chacinska, C. M. Koehler, D. Milenkovic, T. Lithgow, N. Pfanner, Importing mitochondrial proteins: machineries and mechanisms. Cell. 138, 628–644 (2009).

4. A. Sickmann, J. Reinders, Y. Wagner, C. Joppich, R. Zahedi, H. E. Meyer, B. Schönfisch, I. Perschil, A. Chacinska, B. Guiard, P. Rehling, N. Pfanner, C. Meisinger, The proteome of Saccharomyces cerevisiae mitochondria. Proceedings of the National Academy of Sciences. 100, 13207–13212 (2003).

5. R. Ben-Menachem, M. Tal, T. Shadur, O. Pines, A third of the yeast mitochondrial proteome is dual localized: A question of evolution. Proteomics. 11, 4468–4476 (2011).

6. A. Stein, S. Jinks-Robertson, L. Kalifa, E. A. Sia, Members of the RAD52 Epistasis Group Contribute to Mitochondrial Homologous Recombination and Double-Strand Break Repair in Saccharomyces cerevisiae. PLoS Genet. 11, e1005664 (2015).

7. J. Wang, K. Kearney, M. Derby, C. M. Wernette, On the relationship of the ATP-independent, mitochondrial associated DNA topoisomerase of Saccharomyces cerevisiae to the nuclear topoisomerase I. Biochemical and biophysical research communications. 214, 723–729 (1995).

8. J. P. Duxin, B. Dao, P. Martinsson, N. Rajala, L. Guittat, J. L. Campbell, J. N. Spelbrink, S. A. Stewart, Human Dna2 is a nuclear and mitochondrial DNA maintenance protein. Mol Cell Biol. 29, 4274–82 (2009).

9. R. M. Monaghan, A. J. Whitmarsh, Mitochondrial Proteins Moonlighting in the Nucleus. Trends Biochem Sci. 40, 728–735 (2015).

10. M. Dinur-Mills, R. R. Copley, M. Tal, O. Pines, Dual targeted mitochondrial proteins are characterized by lower MTS parameters and total net charge. PLoS One. 3, e2161 (2008).

11. S. A. Ayyub, F. Gao, R. N. Lightowlers, Z. M. Chrzanowska-Lightowlers, Rescuing stalled mammalian mitoribosomes – what can we learn from bacteria? J Cell Sci. 133, jcs231811 (2020).

12. S. D. Moore, R. T. Sauer, The tmRNA system for translational surveillance and ribosome rescue. Annual review of biochemistry. 76, 101–124 (2007).

13. E. Gur, R. T. Sauer, Evolution of the ssrA degradation tag in Mycoplasma: specificity switch to a different protease. Proc Natl Acad Sci U S A. 105, 16113–16118 (2008).

14. D. E. Cameron, J. J. Collins, Tunable protein degradation in bacteria. Nature biotechnology. 32, 1276–1281 (2014).

15. F. D. Bartolomeo, C. Malina, K. Campbell, M. Mormino, J. Fuchs, E. Vorontsov, C. M. Gustafsson, J. Nielsen, Absolute yeast mitochondrial proteome quantification reveals trade-off between biosynthesis and energy generation during diauxic shift. Proc Natl Acad Sci U S A. 117, 7524–7535 (2020).

16. D. Laporte, L. Gouleme, L. Jimenez, I. Khemiri, I. Sagot, Mitochondria reorganization upon proliferation arrest predicts individual yeast cell fate. Elife. 7, 113 (2018).

17. L. E. Bagamery, Q. A. Justman, E. C. Garner, A. W. Murray, A Putative Bet-Hedging Strategy Buffers Budding Yeast against Environmental Instability. Curr Biol (2020), doi:10.1016/j.cub.2020.08.092.

18. L. Galdieri, S. Mehrotra, S. Yu, A. Vancura, Transcriptional Regulation in Yeast during Diauxic Shift and Stationary Phase. Omics J Integr Biology. 14, 629–638 (2010).

19. B. Galeota-Sprung, A. Fernandez, P. Sniegowski, Changes to the mtDNA copy number during yeast culture growth. Roy Soc Open Sci. 9, 211842 (2022).

20. A. Kokina, J. Kibilds, J. Liepins, Adenine auxotrophy – be aware: some effects of adenine auxotrophy in Saccharomyces cerevisiae strain W303-1A. Fems Yeast Res. 14, 697–707 (2014).

21. A. Göke, S. Schrott, A. Mizrak, V. Belyy, C. Osman, P. Walter, Mrx6 regulates mitochondrial DNA copy number in Saccharomyces cerevisiae by engaging the evolutionarily conserved Lon protease Pim1. Mol Biol Cell. 31, 527–545 (2020).

22. R. Jajoo, Y. Jung, D. Huh, M. P. Viana, S. M. Rafelski, M. Springer, J. Paulsson, Accurate concentration control of mitochondria and nucleoids. Science. 351, 169–172 (2016).

23. C. K. Suzuki, K. Suda, N. Wang, G. Schatz, Requirement for the Yeast Gene LON in Intramitochondrial Proteolysis and Maintenance of Respiration. Science. 264, 273–276 (1994).

24. R. Vongsamphanh, P.-K. Fortier, D. Ramotar, Pir1p Mediates Translocation of the Yeast Apn1p Endonuclease into the Mitochondria To Maintain Genomic Stability. Mol Cell Biol. 21, 1647–1655 (2001).

25. X. Cheng, S. Dunaway, A. S. Ivessa, The role of Pif1p, a DNA helicase in Saccharomyces cerevisiae, in maintaining mitochondrial DNA. Mitochondrion. 7, 211–222 (2007).

26. L. D. Cruz-Zaragoza, S. Dennerlein, A. Linden, R. Yousefi, E. Lavdovskaia, A. Aich, R. R. Falk, R. Gomkale, T. Schöndorf, M. T. Bohnsack, R. Richter-Dennerlein, H. Urlaub, P. Rehling, An in vitro system to silence mitochondrial gene expression. Cell. 184, 5824-5837.e15 (2021).

## References cited in Supplementary Materials

27. B. Westermann, W. Neupert, ReferencesMitochondria-targeted green fluorescent proteins: convenient tools for the study of organelle biogenesis in Saccharomyces cerevisiae. Yeast. 16, 1421–1427 (2000).

28. R. D. Gietz, S. Akio, ReferencesNew yeast-Escherichia coli shuttle vectors constructed with in vitro mutagenized yeast genes lacking six-base pair restriction sites. Gene. 74, 527–534 (1988).

29. R. Rothstein, References[19] Targeting, disruption, replacement, and allele rescue: Integrative DNA transformation in yeast. Methods Enzymol. 194, 281–301 (1991).

30. S. Lee, W. A. Lim, K. S. Thorn, ReferencesImproved Blue, Green, and Red Fluorescent Protein Tagging Vectors for S. cerevisiae. Plos One. 8, e67902 (2013).

31. Cold Spring Harb Protoc, in press, doi:10.1101/pdb.rec8585.

32. D. M. MacAlpine, P. S. Perlman, R. A. Butow, ReferencesThe numbers of individual mitochondrial DNA molecules and mitochondrial DNA nucleoids in yeast are co-regulated by the general amino acid control pathway. The EMBO Journal. 19, 767–775 (2000).

33. K. G. Sharma, R. Kaur, A. K. Bachhawat, ReferencesThe glutathione-mediated detoxification pathway in yeast: an analysis using the red pigment that accumulates in certain adenine biosynthetic mutants of yeasts reveals the involvement of novel genes. Arch Microbiol. 180, 108–117 (2003).

34. A. X. Lu, T. Zarin, I. S. Hsu, A. M. Moses, ReferencesYeastSpotter: accurate and parameter-free web segmentation for microscopy images of yeast cells. Bioinformatics. 35, 4525–4527 (2019).

35. J. Schindelin, I. Arganda-Carreras, E. Frise, V. Kaynig, M. Longair, T. Pietzsch, S. Preibisch, C. Rueden, S. Saalfeld, B. Schmid, J.-Y. Tinevez, D. J. White, V. Hartenstein, K. Eliceiri, P. Tomancak, A. Cardona, ReferencesFiji: an open-source platform for biological-image analysis. Nat Methods. 9, 676–682 (2012).

36. R. U. Kulkarni, C. L. Wang, C. R. Bertozzi, ReferencesAnalyzing nested experimental designs—A user-friendly resampling method to determine experimental significance. Plos Comput Biol. 18, e1010061 (2022).

37. A. P. Singh, R. Salvatori, W. Aftab, A. Aufschnaiter, A. Carlström, I. Forne, A. Imhof, M. Ott, ReferencesMolecular Connectivity of Mitochondrial Gene Expression and OXPHOS Biogenesis. Mol Cell. 79, 1051-1065.e10 (2020).

38. S. Gottesman, E. Roche, Y. Zhou, R. T. Sauer, ReferencesThe ClpXP and ClpAP proteases degrade proteins with carboxy-terminal peptide tails added by the SsrA-tagging system. Gene Dev. 12, 1338–1347 (1998).

39. M. Lies, M. R. Maurizi, ReferencesTurnover of Endogenous SsrA-tagged Proteins Mediated by ATP-dependent Proteases in Escherichia coli *. J Biol Chem. 283, 22918–22929 (2008).

40. N. C. Butzin, W. H. Mather, ReferencesCrosstalk between Diverse Synthetic Protein Degradation Tags in Escherichia coli. Acs Synth Biol. 7, 54–62 (2018).

41. U. Teichmann, L. van Dyck, B. Guiard, H. Fischer, R. Glockshuber, W. Neupert, T. Langer, ReferencesSubstitution of PIM1 protease in mitochondria by Escherichia coli Lon protease. J Biol Chem. 271, 10137–10142 (1996).

